# HypoxamicroRNA-210 protects against hepatic steatosis by inhibiting CIDEC expression

**DOI:** 10.64898/2025.12.19.695309

**Authors:** Bo Yan, Sonia Youhanna, Peiyin Chen, Xiuli Jin, Yuma Iwamura, Aurino Kemas, Jacob Grünler, Sampath Narayanan, Allan Zhao, Qiaolin Deng, Norio Suzuki, Yiling Li, Volker Martin Lauschke, Xiaowei Zheng, Sergiu-Bogdan Catrina

## Abstract

**Background and Aims:** Metabolic dysfunction-associated steatotic liver disease (MASLD) is a major global health burden. Although hypoxia is known to contribute to MASLD pathogenesis, the role of hypoxia signaling remains poorly defined. We investigated whether HypoxamicroRNA-210 (miR-210), a key hypoxia-inducible microRNA, regulates hepatic lipid metabolism and MASLD development.

**Methods:** Serum miR-210 levels were quantified in MASLD patients and matched controls. Human hepatic spheroids and HepG2 cells were exposed to fatty acids to assess miR-210 induction and lipid accumulation. miR-210 knockout mice were fed a Western diet to evaluate hepatic steatosis and transcriptomic changes using RNA sequencing. RNA pull-down and 3’UTR-driven luciferase reporter assays were employed to identify miR-210 targets. Functional effects of miR-210 mimic were examined in knockout mice, db/db mice, and *in vitro* human hepatic spheroid models.

**Results:** Serum miR-210 levels were significantly reduced in MASLD patients compared with matched controls. Consistently, human hepatic spheroids did not appropriately increase miR-210 expression in response to fatty acid-induced intracellular hypoxia. This blunted miR-210 response contributed to hepatic lipid accumulation, as loss of miR-210 in a mouse model of MASLD led to increased hepatic lipid deposition and activation of lipid metabolic pathways. We identified CIDEC as a direct miR-210 target mediating its inhibitory effects on hepatic lipid accumulation, and restoring miR-210 expression suppressed CIDEC and reduced hepatic lipid content in knockout mice on a Western diet. Moreover, miR-210 attenuated lipid accumulation in both *in vitro* human hepatic spheroids and *in vivo* db/db mice models of MASLD.

**Conclusions:** miR-210 protects against hepatic steatosis by inhibiting CIDEC expression, suggesting miR-210-CIDEC axis as a promising therapeutic target for reducing hepatic lipid accumulation and preventing MASLD progression.

**Impact and Implications:** This study addresses a critical gap in understanding how hypoxia signaling shapes MASLD and uncovers a novel pathogenic mechanism of hepatic steatosis arising from fatty acids-induced dysregulation of miR-210. Impaired hypoxia responses, via blunted miR-210 induction, contribute to hepatic lipid accumulation through upregulation of CIDEC, a newly identified target of miR-210. These findings establish miR-210 as a novel regulator of hepatic lipid homeostasis and underscore its therapeutic potential. Interventions aimed at restoring miR-210 function in the liver may offer a promising strategy to ameliorate hepatic steatosis and prevent MASLD progression.

## Introduction

Metabolic dysfunction-associated steatotic liver disease (MASLD) has emerged as the most prevalent chronic liver disorder globally, affecting an estimated 38 % of the adult population ^[1]^. MASLD is a progressive disease characterized by excessive hepatic lipid accumulation in the absence of significant alcohol consumption. It encompasses a spectrum of histological phenotypes, beginning with hepatic steatosis and potentially progressing to metabolic dysfunction-associated steatohepatitis (MASH), cirrhosis, and hepatocellular carcinoma ^[2]^. As a major driver of liver-related morbidity and mortality, MASLD is also an independent risk factor for cardiovascular events ^[3]^ and chronic kidney disease ^[4]^. Currently, therapeutic options for MASLD are limited, primarily relying on lifestyle modifications, with few pharmacological therapies available ^[5]^.

The hallmark of MASLD is hepatic steatosis, defined as the non-physiological accumulation of triglycerides (TG) in lipid droplets in hepatocytes ^[1]^. Once considered passive lipid storage depots, lipid droplets are now recognized as dynamic organelles regulating lipid metabolism, transport, and signaling. These functions are largely governed by proteins residing on the lipid droplet surface, which regulate the metabolic fate of stored lipids ^[6, 7]^. Among these, the CIDE (cell death–inducing DNA fragmentation factor-α–like effector) family proteins has been the focus of extensive investigation ^[8, 9]^.

MASLD pathogenesis involves complex interactions between hepatic lipid accumulation, inflammation, oxidative stress, endoplasmic reticulum stress, and mitochondrial dysfunction ^[10]^. Hepatic hypoxia has emerged as a significant contributor to MASLD ^[11],^ arising from impaired sinusoidal perfusion as lipid infiltration progresses ^[12]^ and increased oxygen consumption for beta oxidation of the excessive fatty acids (FA) ^[13]^. Notably, high-fat diet feeding can induce hepatic hypoxia within seven days ^[14, 15]^. However, the underlying mechanisms of hypoxia in the progress of MASLD remain poorly defined.

Cellular adaptation to oxygen deficiency is primarily mediated by hypoxia-inducible factors (HIFs), which orchestrate the transcriptional activation of hundreds of genes to sustain homeostasis under hypoxic conditions ^[16]^. However, the contribution of HIFs to MASLD remains debated ^[11, 17],^ likely influenced by isoform specificity (HIF-1 vs HIF-2) and disease stage. Moreover, their expansive regulatory network adds significant complexity to downstream effects, particularly in the multifactorial context of MASLD. In contrast, hypoxia-responsive microRNAs (HypoxamicroRNAs), notably miR-210, represent more selective and streamlined mediators of hypoxic signaling ^[18, 19]^. As a robust HIF-1 target, miR-210 exerts critical modulatory effects on mitochondrial metabolism ^[20, 21]^ and lipid homeostasis ^[22]^.

Understanding how miR-210 integrates hypoxia signaling with hepatic lipid metabolism could provide novel insights into MASLD pathogenesis. Moreover, deciphering its downstream targets may identify new therapeutic opportunities aimed at modulating maladaptive responses to chronic metabolic and hypoxic stress in the liver.

We therefore investigated the role of miR-210 in the pathogenesis of MASLD. We found that serum miR-210 levels were reduced in MASLD patients. Mechanistically, miR-210 expression failed to increase in response to FA-induced intracellular hypoxia in human hepatic spheroids. The direct contribution of miR-210 dysregulation to MASLD pathogenesis was confirmed in miR-210 knockout mice fed a Western Diet (WeD), which exhibited increased hepatic lipid accumulation that was reversed by miR-210 reconstitution. The lipid droplet-associated protein CIDEC, also known as FSP27 (fat-specific protein 27), was identified as a direct target of miR-210 and a key mediator of its effects on hepatic lipid accumulation. Moreover, administration of miR-210 mimic mitigated hepatic steatosis in both human hepatic spheroids and db/db mice, well-established models of MASLD.

## Methods

### Human serum samples

A total of 21 subjects with MASLD and 11 healthy, age-matched controls were enrolled at the First Affiliated Hospital of China Medical University, as illustrated in the flowchart in Fig. S1. Demographic and clinical characteristics of the study participants are summarized in Table S1. The clinical study was approved by the Ethics Committee of China Medical University (Ethics Approval No. 2024-577) and conducted in accordance with the principles of the Declaration of Helsinki. Written informed consent was obtained from all participants.

Healthy subjects without hepatic steatosis were identified by abdominal CT or ultrasound. All healthy controls had no evidence of hyperlipidemia and had body mass index (BMI) below 30 kg/m^2^. Subjects with MASLD were diagnosed according to the MASLD diagnostic criteria ^[23],^ which require evidence of hepatic steatosis (confirmed by Fibrotouch that Fatty decay > 244 dB/m) along with at least one of the following five cardiometabolic risk factors: (1) BMI ≥ 23 kg/m^2^ (Asian criteria) or wrist circumstance > 94 cm (men) or > 80cm (women); (2) Fasting serum glucose ≥ 5.6 mmol/L or 2-hour post-load glucose levels ≥ 7.8 mmol/L or HbA1c ≥ 5.7% or type 2 diabetes (T2D) or treatment for T2D; (3) Blood pressure ≥ 130/85 mmHg or specific antihypertensive drug treatment; (4) Plasma triglycerides ≥ 1.7mmol/L or lipid lowering treatment; (5) Plasma HDL-cholesterol ≤ 1.0mmol/L (men) and ≤ 1.3mmol/L (women) or lipid lowering treatment. Exclusion criteria included a history of smoking or excessive alcoholic consumption; the presence of acute (e.g., infections) or chronic diseases (e.g., inflammatory bowel disease, malignancies); liver dysfunction (ALP ≥ 125 U/L or ALT and AST levels over twice the upper limit) attributed to viral, drug-induced, autoimmune, or unidentified causes; and the use of hepatoxic or lipid-lowering medications.

Whole blood of enrolled clinical subjects was collected via venipuncture and allowed to clot at room temperature for 30 min in plain collection tubes. Serum was then separated after centrifugation at 2000 × g for 10 min. Analysis of metabolic and other biological parameters in Table S1 were performed by the Clinical Chemistry Laboratory at the First Affiliated Hospital of China Medical University. An aliquot of 500 μL of serum was collected and stored at -80 °C for further analysis.

### Analysis of NCBI dataset GSE185062

To examine an additional clinical cohort of MASLD, the GEO database (https://www.ncbi.nlm.nih.gov/geo/) was searched using the keywords “NAFLD”, “microRNA” and “human”. Dataset GSE185062 ^[24],^ comprising serum non-coding RNA sequencing data from 183 patients with biopsy-confirmed hepatic steatosis and 10 healthy controls, was selected for analysis. Raw counts were normalized to counts per million (CPM), and miR-210-3p CPM values were extracted and compared with healthy controls both collectively and across patient groups stratified by disease severity according to the categories defined in the original publication ^[24]^.

### Animal models

miR-210 KO was previously described ^[25]^ and locally validated ^[26]^. Twelve-week-old male miR-210 knockout (KO) and littermate wild-type (WT) male mice were randomly assigned to receive either a standard Control Diet (CoD) or WeD (RD western diet, D12079Bi) for 10 weeks. BKS(D)-*Lepr^db^*/JOrIRj (db/db) male mice were obtained from Janvier Labs. The mice were housed up to five per cage at 24 ± 2 °C, in a 12-hour light/dark cycle, and had access to CoD or WeD and water *ad libitum*. The mice were randomized into groups receiving control microRNA mimic or miR-210 mimic according to their body weight and blood glucose levels. The experimental animal procedure was approved by the North Stockholm Ethical Committee for the Care and Use of Laboratory Animals.

### MicroRNA mimic injection and tissue collection

Custom-stabilized miRIDIAN microRNA-210-3p mimic (miR-210 mimic, C-310570-5) and miRIDIAN microRNA Mimic Negative Control #1 (control mimic, CN-001000-01) were purchased from Horizon Discovery. The microRNA mimic was packed into phospholipid-oil emulsion as previously described ^[20],^ by passing the mixture of microRNA mimic and MaxSuppressor^TM^ In Vivo RNA-LANCEr II agent (BIOO Scientific, USA) through the Lipid Extruder (BIOO Scientific, USA). 0.5 μg/g miR-210 mimic or control mimic were delivered to each mouse by tail vein injection, and the mice were sacrificed five days thereafter. Liver tissues were collected, snap frozen in liquid nitrogen, and stored at -80 ℃, or preserved in RNAlater solution (ThermoFisher Scientific, AM7024) overnight and subsequently stored at - 20℃.

### HepG2 cell culture

Human hepatoma cell line HepG2 cells (ATCC, USA) were cultured in Dulbecco’s modified Eagle’s medium (DMEM, 4.5 g/L glucose) supplemented with 100 U/ml penicillin, 100 μg/mL streptomycin, and 10% FBS (ThermoFisher Scientific). All cells were confirmed to be mycoplasma-free using the MycoAlert PLUS Mycoplasma Detection Kit (LONZA). Cells were maintained in a humidified incubator at 37 °C with 5% CO_2_. To emulate steatosis, HepG2 cells were exposed to 400 μM palmitic acids for 24 hours with the final 16 hours conducted under hypoxic conditions (1% O_2_) in Hypoxia Workstation INVIVO2 (Ruskinn) prior to harvest.

To investigate the effects of miR-210, HepG2 cells were transfected with 1nM control mimic or miR-210 mimic using Lipofectamine RNAiMAX (ThermoFisher Scientific) according to the manufacturer’s protocol. CIDEC overexpression was achieved by co-transfection with 1 μg FLAG-CIDEC-GFP plasmid or control FLAG-GFP vector (both purchased from MiaoLingPlasmid) using Lipofectamine 3000 (Invitrogen, L3000015). After 24 hours of transfection, cells were treated with 400 μM palmitic acid (Sigma, P0500) conjugated to fatty acid–free bovine serum albumin (BSA, 10%) for 24 hr. Control cells received 10% BSA alone. For hypoxia treatment, cells were placed in a Hypoxia Workstation INVIVO2 (Ruskinn) (1% O₂) during the last 16 hours of experiments. After treatment, cells were harvested for total RNA and protein extraction, triglyceride quantification, Oil Red O staining, and confocal microscopy.

### Hepatic spheroids culture

Cryopreserved primary human hepatocytes (BioIVT) were cultured in 96-well ultra-low-attachment plates (Corning) using spheroid culture medium (Williams’ medium E containing 11 mM glucose and supplemented with 2 mM L-glutamine, 100 U/mL penicillin, 100 µg/mL streptomycin, 100 pM insulin, 5.5 µg/mL transferrin, 6.7 ng/mL sodium selenite, 100 nM dexamethasone) supplemented with 10% heat-inactivated fetal bovine serum (FBS), as previously described ^[27, 28]^. After spheroids had formed, FBS was phased out before further exposures. The cells were transfected during seeding with 1 nM control mimic or miR-210 mimic using RNAiMax (ThermoFisher Scientific) according to the manufacturer’s protocol. From day 7 post-seeding, spheroids were exposed to either physiological medium (1720 pM insulin) or lipogenic medium containing 1.7 µM insulin, 240 μM palmitic acid and 240 μM oleic acid, mimicking human plasma levels of free fatty acids. Medium was exchanged every 48–72 hours over a 7-day period. On the sixth day of lipogenic exposure, spheroids were incubated under normoxic (21% O_2_) or hypoxic (1% O_2_) conditions in a Hypoxia Workstation INVIVO2 (Ruskinn) for 24 hours. For hypoxia evaluation, spheroids were treated with 200 μM pimonidazole for 2 hours prior to collection.

### Dual-luciferase reporter assay

The WT human *CIDEC* 3′UTR or a mutant 3′UTR with deletion of the predicted miR-210 binding site was cloned downstream of the Renilla luciferase gene in the psiCHECK-2 vector (Promega). HepG2 cells were seeded in 12-well plates and co-transfected with 100 ng of the WT or mutant reporter plasmids and 5 nM miR-210 mimic or control mimic using Lipofectamine 3000 and RNAiMAX according to the manufacturer’s instructions. After 24 hours of transfection, cells were exposed to 400 μM palmitic acid and hypoxia for additional 24 hours before harvesting. Luciferase activities were measured using the Dual-Luciferase Reporter Assay System (Promega). Renilla luciferase activity was normalized to firefly luciferase activity from the same vector to control for transfection efficiency. Relative luciferase activity was expressed as fold induction relative to cells transfected with WT reporter and control mimic.

### miRNA-mRNA complex pulldown

To validate *CIDEC* as a target gene of miR-210, an optimized RNA pulldown protocol was adapted from previous reports ^[29–32]^. HepG2 cells were transfected with 20nM biotinylated control mimic or miR-210 mimic (Qiagen, 339178) using Lipofectamine RNAiMAX. Cells were treated with 400 μM palmitic acids for 24 hours and exposed to hypoxia for the final 16 hours.

Prior to collection, cells were crosslinked with 1% paraformaldehyde (PFA) for 10 minutes, followed by quenching with 1.25 M glycine. Cell pellets were lysed in ice-cold cell lysis buffer (150 mM NaCl, 25 mM Tris-HCl pH-7.5, 5mM DTT, 0.5% IGEPAL, 60 U/mL RNase inhibitor, and 1x Protease Inhibitor). The lysate was adjusted to a final NaCl concentration of 1M. 50μL lysate was reserved as input.

Streptavidin-coated magnetic beads (Cytiva, 65152105050250) were pre-blocked overnight in blocking buffer (1 µg/µL BSA, 2 µg/µL Yeast tRNA) and resuspended in pull-down wash buffer (10 mM KCl, 1.5 mM MgCl2, 10 mM Tris-Cl pH 7.5, 5 mM DTT, 1 M NaCl, 0.5% IGEPAL, 60 U/mL RNase Inhibitor, and 1x Protease Inhibitor). Cell lysates were incubated with the blocked beads for 1.5 hours at room temperature, followed by three washes with pull-down wash buffer. Protein degradation was carried out by incubating the pulldown and input samples with proteinase K (1 mg/mL) for 45 min at 50 °C then denatured for 10 min at 95 °C. Beads were then separated from the supernatant using magnetic stand. Total RNA was extracted from the pulldown and input samples and followed by qRT-PCR analysis.

### RNA purification

Serum samples were briefly warmed to 37℃ and then completely thawed at room temperature. The thawed serum samples were centrifuged at 3000 × g for 15 minutes at 4 ℃. A 200 uL aliquot of the clarified serum was used for RNA extraction. Total RNA, including microRNA, was purified using the miRNeasy Serum / Plasma Advanced Kit (Qiagen, 217204).

Liver tissue samples (20-30 mg) preserved in RNAlater solution were used for RNA isolation. Total RNA, including microRNAs, was extracted using Nucleospin miRNA kit (Techtum, 740971). Total RNA from hepatic spheroids and HepG2 cells were extracted using Direct-zol^TM^ RNA Microprep kit (Zymo Research, R2060) and Direct-zol^TM^ RNA Miniprep kit (Zymo Research, R2050), respectively. All RNA purification procedures were performed in accordance with the manufacturer’s protocols. Genomic DNA was eliminated using DNase during the procedure. The concentration and purity of extracted RNA were assessed using a NanoDrop spectrophotometer (ThermoFisher Scientific).

### Primer Design for SYBR Green qPCR

Primers were designed to amplify target genes using SYBR Green chemistry, following MIQE guidelines ^[33]^. Sequences were selected using Primer-BLAST (NCBI) to ensure specificity and avoid secondary structures. Amplicon lengths were restricted to 80–200 bp to optimize amplification efficiency. Primers were designed to span exon–exon junctions where possible to minimize genomic DNA amplification. GC content was maintained between 40–60%, and melting temperatures (Tm) were set between 58–62°C with minimal difference (<1 °C) between forward and reverse primers. Specificity was confirmed by in silico BLAST analysis against the reference genome. All primers were synthesized by ThermoFisher Scientific and validated experimentally by melt curve analysis to exclude primer dimer and to confirm single amplicon formation. Amplification efficiency was determined using standard curves generated from serial dilutions of cDNA, and only primers with efficiencies between 90–110% and R² > .99 were used for subsequent analyses. The primer sequences are provided in the Supplementary Methods.

### Reverse transcription and quantitative RT-PCR

For mRNA detection, cDNA was synthesized using the High-Capacity cDNA Reverse Transcription Kit (ThermoFisher Scientific, 4368814). Quantitative RT-PCR (qRT-PCR) was performed using the QuantStudio 7 Pro Real-Time PCR System (Applied Biosystems) with the Powerup SYBR Green Master Mix (ThermoFisher Scientific, A25742) according to the manufacturer’s protocols. The internal control for mRNA expression was *ACTB*.

For the detection of miR-210 expression in mouse liver tissue, human hepatic spheroids and HepG2 cells, cDNA was synthesized using the TaqMan Advanced miRNA cDNA Synthesis Kit (ThermoFisher Scientific, A28007). qRT-PCR was performed using the Taqman Fast Advanced Master mix (ThermoFisher Scientific, A44360) and TaqMan Advanced miRNA assays hsa-miR-210-3p (ThermoFisher Scientific, 477970) and hsa-miR-16-5p (ThermoFisher Scientific, 47786). To assess the expression of miR210 in serum samples, cDNA was synthesized using the miRcute Plus miRNA First-Strand cDNA kit (Tiangen Biotech, KR211). qRT-PCR was performed on a 7500 FAST Real-Time PCR System (Applied Biosystems) using miRcute Plus miRNA qPCR Kit (SYBR Green) (Tiangen Biotech, FP411). miR-16 served as the internal reference control for miR-210 expression.

### RNA sequencing and data analysis

Total RNA from liver tissue was used to construct bulk RNA sequencing libraries following the prime-seq protocol, as previously described ^[34]^. cDNA and final library concentrations were measured using the Qubit 1X dsDNA High Sensitivity Assay Kit (ThermoFisher Scientific), while size distribution and quality were assessed using the Bioanalyzer High Sensitivity DNA Kit (Agilent Technologies) according to the manufacturers’ instructions. Libraries were sequenced on an Illumina NovaSeq 6000 platform at Novogene Ltd, generating an average of 20 million reads per sample.

Raw sequencing data were processed in R Studio (v22.0.3) using the *Seurat* package (version 4.3.0) ^[35]^. Quality control included assessments of read counts, number of detected genes and mitochondrial gene content. All samples met quality criteria. Mitochondrial genes and lowly expressed genes (total count ≤ 20 across all samples) were excluded. Data were normalized using lognormalization, in which counts were divided by the total counts per sample, scaled by 10,000, and log-transformed.

Gene Ontology (GO), KEGG and Hallmark gene set enrichment analysis (GSEA) ^[36]^ of all the detected genes were performed using gene sets from the Molecular Signatures Database (MSigDB, version 2025.1) and the *clusterProfiler* ^[37]^ and *ggplot2* packages ^[38]^. Gene sets with a normalized enrichment score (NES) ≥ 1 and q value < 0.05 were considered significantly activated, whereas those with NES ≤ -1 and q value < 0.05 were considered significantly downregulated.

Differentially expressed genes (DEGs) were identified using the *DESeq2* package (version 1.40.2) ^[39]^. Genes with |Log_2_ (fold change) (FC)| > 0.5 and adjusted *p*-value (*P*adj) < 0.1 were considered differentially expressed. Volcano plots were generated with *EnhancedVolcano*, and Venn diagrams with *Venn*. GO pathway enrichment analysis of DEGs was performed using *clusterProfiler* package and visualized with *ggplot2* package.

### Protein extraction and Western blotting analysis

Mouse liver tissue, hepatic spheroids and HepG2 cell pellets were homogenized in pre-cooled RIPA buffer containing proteinase and phosphatase inhibitors (ThermoFisher Scientific) using TissueLyser II (Qiagen). Lysates were prepared by centrifugation at 20,000 × g for 30 minutes at 4 °C. Protein concentration were measured using Bicinchoninic Acid Protein Assay kit (Sigma, B9643) and separated by SDS-PAGE (Genescript, M00656) and blotted onto PVDF membranes (BioRAD, 162-0260). Membranes were blocked in Intercept Blocking Buffer (LI-COR, 927-60001) for 1 hour at room temperature, followed by overnight incubation at 4 °C with primary antibodies diluted in Intercept Blocking Buffer containing 0.2% Tween20. The following primary antibodies were used: rabbit anti-pimonidazole (1:1000, Hypoxyprobe, PAb2627AP), rabbit anti-CIDEC (1:500, Abcam, ab198204), mouse anti-GPD1 (1:100, Santa Cruz, sc-376219), rabbit anti-β-actin (1:5000, Abcam, ab8227). After three washes (3 minutes each) with TBS containing 0.1% Tween20 (TBS-T), membranes were incubated with IRDye 800 goat anti-rabbit, IRDye 680 goat anti-rabbit, or IRDye 800 goat anti-mouse secondary antibodies (1:20000, LI-COR). After three washes (10 minutes each) with TBS-T, membranes were scanned using Odyssey Clx Imaging System (LI-COR). Quantification of western blots were performed using ImageJ (Version 1.53).

### Histology

Approximate 30 mg frozen liver tissues were embedded in OCT embedding compound (Tissue-Tek, 4853) and stored at -80 ℃ until use. During cryosection, samples were sectioned into 8 μm slices using an N35 microtome blade (Pfm medical, 207500006). Sections were air-dried at room temperature for 1 hour, fixed in 4% formaldehyde for 10 minutes, and rinsed once in PBS. The slides were then left at room temperature overnight before being stored at - 20℃. Frozen sections (8 μm thickness) were rinsed in 1× PBS for 3 minutes to remove OCT embedding compounds. After a 1-minute wash under running tap water, slides were immersed in hematoxylin (Histolab, 01820) for 10 minutes, followed by a 6-minute rinse under running tap water. Sections were then stained with eosin (Histolab, 01650) for 30 seconds and rinsed again with running tap water for 30 seconds. Slides were dehydrated through a graded ethanol series (70%, 95%, 100%) and cleared in xylene (Histolab, 02070), then mounted using limonene mounting medium (Abcam, ab104141). Images were captured using a bright-field microscope (Leica, DM3000).

Histological assessment was performed on images at 20× magnification using the NAFLD Activity Score (NAS) system ^[40]^. The NAS ranges from 0 to 8 and is calculated as the sum of scores for steatosis (0-3), lobular inflammation (0-3) and hepatocyte ballooning (0-2).

### Oil Red O staining

Oil-red O stock solution was prepared by dissolving 0.3g Oil-red O power (Sigma, O0625) in 100 mL 100% isopropanol. For staining, Oil-red O stock solution was diluted in MQ water at a 3:2 ratio and filtered before use. Frozen liver sections (8 μm thickness) were air-dried at room temperature for more than 30 minutes, then briefly dipped twice in 60% isopropanol (3 sec each). Sections were incubated with the oil-red O working solution in dark for 15 min, washed three times in 60% isopropanol, and rinsed twice with MQ water. For bright-field images, slides were counterstained with hematoxylin for 3min and rinsed twice with MQ water. Sections were mounted using DAKO aqueous mounting medium and imaged under a bright-field microscope (Leica, DM3000). Stained lipid area in 20× magnification images were quantified using ImageJ software. For confocal microscopy, nuclei were counterstained with DAPI (1 μg/mL; ThermoFisher Scientific) and mounted using Fluorescence Mounting Medium (Dako, Cat# S3023). Images were acquired with the LSM900-Airy Confocal Microscope (Zeiss, Germany).

### Measurement of triglyceride content

Liver tissue and HepG2 cells were homogenized in 5% IGEPAL (Sigma, 56741). Lysates were heated to 90℃ for 3min twice and followed by centrifugation at 4℃ for 30 min. Supernatant was 1:10 diluted and assayed for triglycerides using the Triglyceride Quantification Kit (Sigma, MAK266) according to the manufacturer’s instructions. The absorbance was measured at 570 nm using a microplate reader, and triglyceride levels were normalized to protein concentrations.

Quantification of triglyceride in human hepatic spheroid samples was performed with either Triglyceride Quantification Kit (Sigma, MAK266) where 48 spheroids were homogenized in 60uL 5% IGEPAL, or with AdipoRed assay (Lonza, PT-7009) following the manufacturer’s protocols. DNA concentration, determined using Quant-iT dsDNA assay kit (ThermoFisher Scientific, Q33120), was used for normalization.

### Statistical Analysis

All statistical analyses were based on independent biological replicates. Statistical analysis was conducted using GraphPad Prism software. Normality was analyzed using Shapiro-Wilk test. Comparisons between two groups were assessed using an unpaired two-tailed Student’s t-test. For comparisons involving two independent variables, two-way ANOVA was performed, followed by Bonferroni post hoc tests. Correlation analysis was performed using Pearson correlation test. When data were not normally distributed, the Mann-Whitney U test was used for comparisons between two groups, and the Kruskal-Wallis test for comparisons among more than two groups. A *p*-value < 0.05 was considered statistically significant. Data are presented as mean ± standard error of the mean (SEM).

## Results

### Reduced serum levels of miR-210 in subjects with MASLD

We first analyzed miR-210 levels in serum from MASLD patients and in healthy subjects. Serum miR-210 levels were significantly reduced in subjects with MASLD. Notably, serum miR-210 levels negatively correlated with triglyceride concentrations and positively correlated with HDL levels (Fig. 1A-1D and S1, Table S1-S2), suggesting a potential link between miR-210 and dysregulated lipid metabolism in MASLD.

**Fig. 1.**
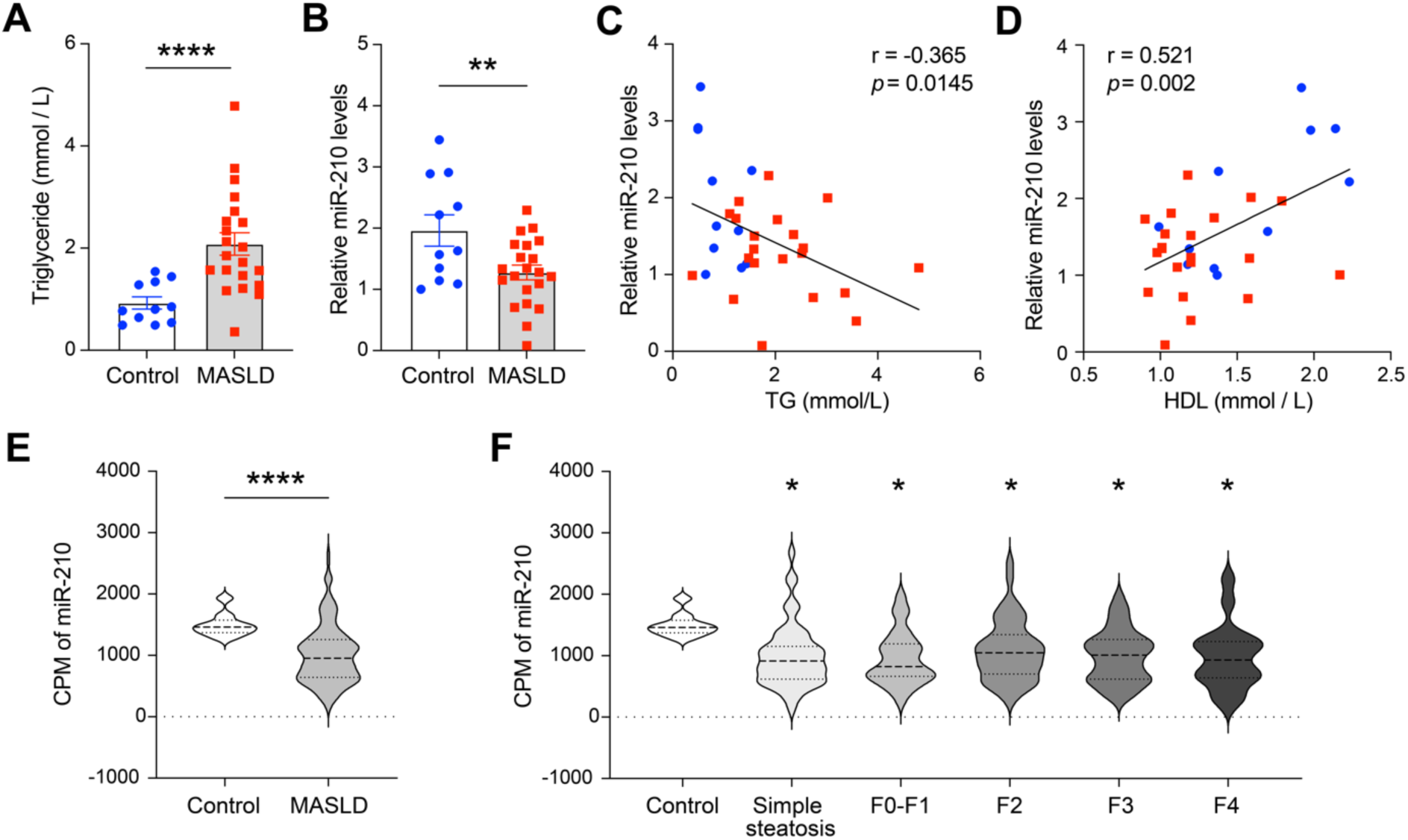
Serum miR-210 expression in MASLD patients and its correlation with lipid profile. (A-B) Serum triglyceride (TG) (A) and miR-210 (B) levels from MASLD patients (n=21) and healthy controls (n=11). **(C-D)** Correlation analysis between serum miR-210 levels with TG (C) and High-density lipoprotein (HDL) cholesterol (D) (n=32). Data are shown as mean ± SEM. **(E)** Counts per million total reads (CPM) of miR-210 in serum of MASLD patients of all the stages (n=183) were lower than healthy control subjects (n=10) in NCBI dataset GSE185062. **(F)** CPM of miR-210 were lower in serum of MASLD patients of the indicated stages than healthy control subjects in NCBI dataset GSE185062. Control (n=10), Simple steatosis (n=55), F0-F1 (n=25), F2 (n=47), F3 (n=38), F4 (n=18). Statistical significance was determined using unpaired Student’s t-test (A-B), Pearson correlation test (C-D), Mann-Whitney U test (E) and Kruskal-Wallis test (F). \**p* < 0.05; \*\**p* < 0.01; \*\*\*\**p* < 0.0001.

To further validate this finding, we analyzed *NCBI dataset GSE185062* which includes non-coding RNA sequencing data from the serum of 183 individuals with biopsy-confirmed hepatic steatosis and 10 healthy controls (Fig. S2) ^[24]^. Consistent with our results, miR-210 counts were significantly lower in patients with MASLD across all stages, including simple steatosis without inflammation or fibrosis, the early stages without fibrosis (F0) or with negligible fibrosis (F1), and even in later stages with advanced fibrosis (F2), bridging fibrosis (F3) and cirrhosis (F4) (Fig. 1E-1F). Collectively, these results establish that serum miR-210 levels are reduced in MASLD patients.

### Fatty acids suppress miR-210 induction by hypoxia in human hepatic spheroids

To investigate how FA regulates miR-210 in the liver, we treated human hepatic spheroids with FA. FA treatment induced intracellular hypoxia in human hepatic spheroids, as demonstrated by increased levels of pimonidazole adducts (Fig. 2A), an established hypoxia marker ^[41–43]^. FA treatment further enhanced pimonidazole adduct formation in spheroids cultured under hypoxic conditions. Since miR-210 is a well-established HypoxamicroRNA, we anticipated higher miR-210 expression in spheroids exposed to FA. As expected, hypoxia alone induced miR-210; however, the additional hypoxia triggered by FA did not lead to further increase in miR-210 expression in hypoxia (Fig. 2B). These findings indicate that, although FA induces hypoxia in human hepatic spheroids, it paradoxically suppresses miR-210 induction. Indeed, exposure to higher FA concentrations in hypoxia further reduced miR-210 levels (Fig. 2C), supporting an inhibitory effect of FA on miR-210 expression.

**Fig. 2.**
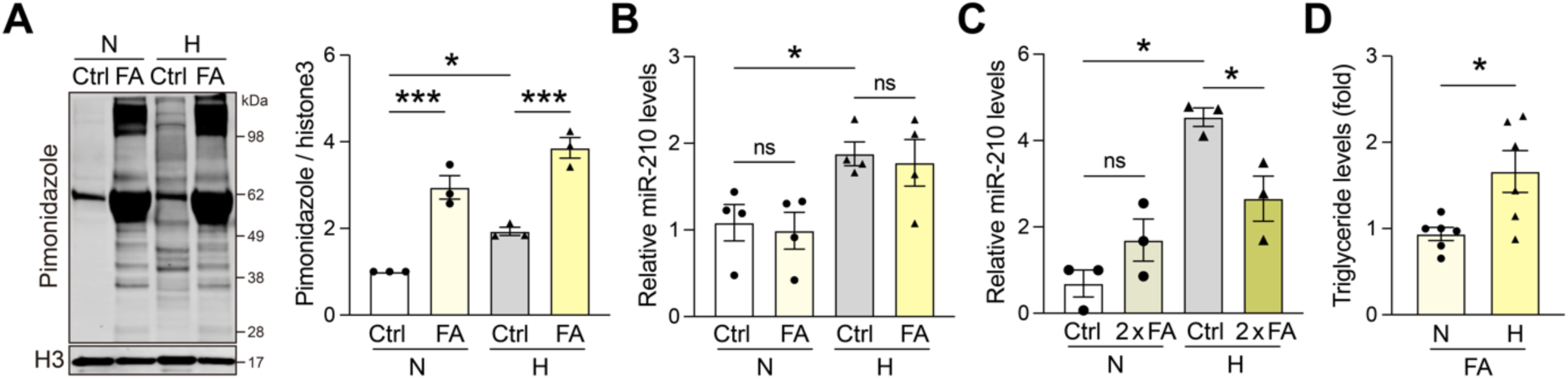
Fatty acids induce hypoxia but blunt miR-210 induction in human hepatic spheroids. Human hepatic spheroids were treated with fatty acids (FA) or vehicle (Ctrl) under normoxic (N) or hypoxic (H) conditions. **(A)** Representative images and quantification of Western blotting results showing pimonidazole adduct levels in the human hepatic spheroids (n=3). Histone H3 (H3) was used as a loading control. **(B-C)** Dose-dependent suppression of miR-210 by fatty acids in human hepatic spheroids. Relative miR-210 expression in human hepatic spheroids exposed to FA (B) or twice more concentrated FA (2 x FA) (C) (n=3-4). **(D)** Triglyceride levels in FA-treated human hepatic spheroids under normoxic or hypoxic conditions (n=5). Data are presented as mean ± SEMs. Statistical significance was determined using Two-way ANOVA with Bonferroni post hoc test (A-C) and unpaired Student’s t-test (D). \**p* < 0.05; \*\*\**p* < 0.001; ns: not significant.

Interestingly, hypoxia was also followed by increased TG accumulation in the human hepatic spheroids (Fig. 2D), which may result from insufficient miR-210 induction in hypoxia due to the inhibition by FA.

### MiR-210 deficiency promotes hepatic lipid accumulation in MASLD mouse model

To assess the role of miR-210 in hepatic lipid accumulation *in vivo*, wild-type (WT) and miR-210 knockout (KO) mice were fed either a WeD or a CoD (Fig. 3A - 3B). Compared to WT mice, miR-210 KO mice on WeD exhibited an accelerated body weight gain (Fig. 3C). Histological and biochemical analyses revealed that miR-210-KO mice on WeD developed more severe hepatic steatosis, as indicated by higher NAFLD Activity Score (NAS), increased hepatic lipid deposition, and elevated TG levels in the liver (Fig. 3D-3F).

**Fig. 3.**
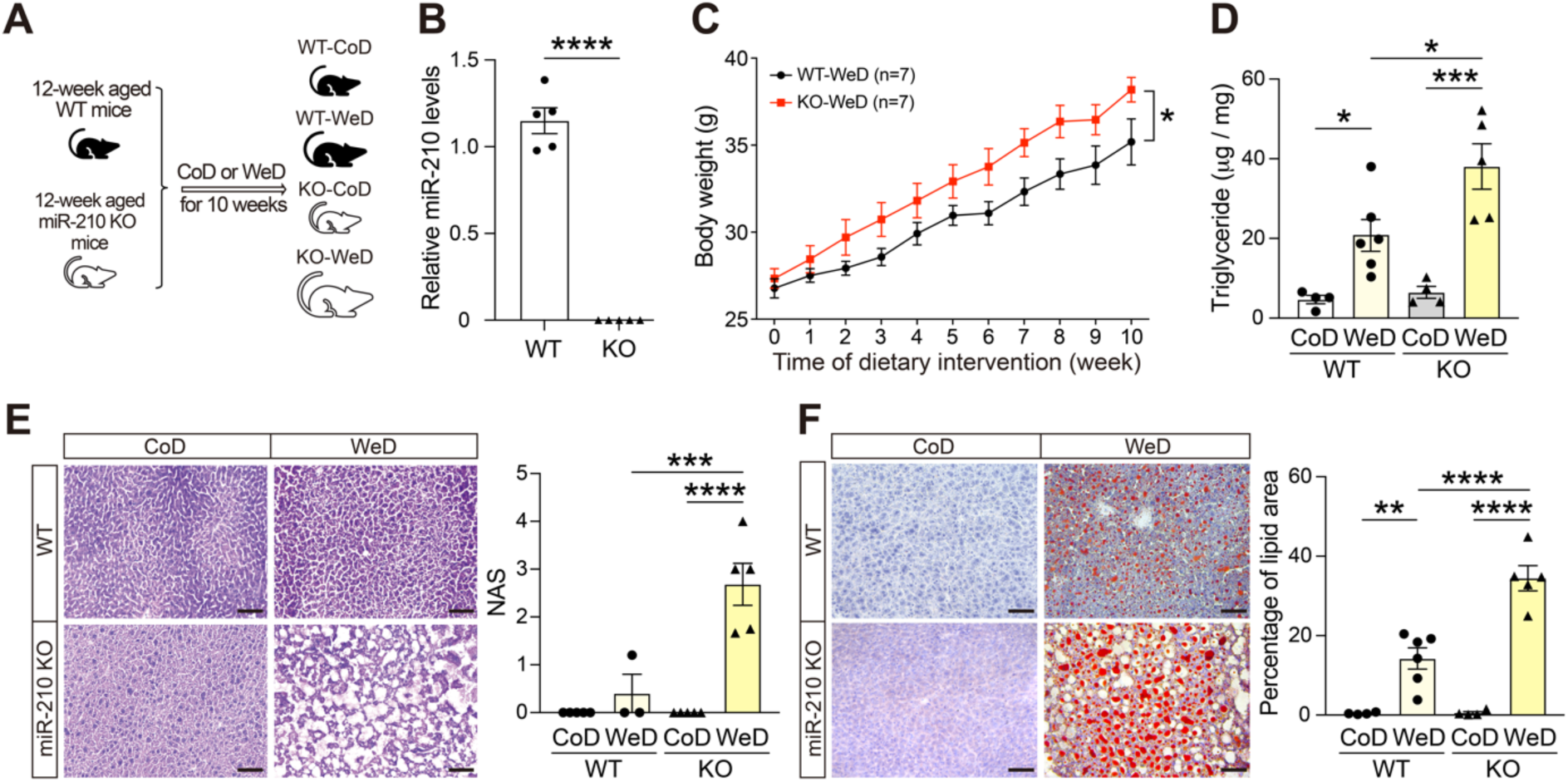
miR-210 deficiency promotes hepatic lipid accumulation in a mouse model of MASLD. Wild-type (WT) and miR-210 knockout (KO) mice were fed a Control (CoD) or Western Diet (WeD) for 10 weeks (n=5-7). **(A)** Schematic experimental design. **(B)** Relative miR-210 expression in the livers of WT and miR-210 KO mice. **(C)** Body weight curves for WeD-fed WT and miR-210 KO mice. **(D)** Hepatic triglyceride (TG) concentrations. **(E)** Histological analysis (H&E staining) of liver sections and quantification of NAFLD Activity Score (NAS). **(F)** Oil Red O staining of hepatic lipid deposition. Scale bar: 50μm. Data are presented as mean ± SEM. Statistical significance was determined using unpaired Student’s t-test (B) and Two-way ANOVA with Bonferroni post hoc test (C-F). \**p* < 0.05; \*\**p* < 0.01; \*\*\**p* < 0.001; \*\*\*\**p* < 0.0001.

Gene expression analysis using qRT-PCR further demonstrated upregulation of lipid metabolism-related genes, such as CD36 and SREBF1, in the livers of miR-210 KO mice fed the WeD. In contrast, expression of inflammation markers (*F4/80* and *IL1B*) and fibrosis-related genes (*ACTA2*, *TGFB*, and *MMP9*) remained unchanged, suggesting that inflammation and fibrosis have not been affected by miR-210 KO (Fig. S3).

These findings, consistent with data from patients and human hepatic spheroids, suggest that miR-210 plays a protective role against hepatic lipid accumulation in MASLD, likely through the regulation of lipid metabolism.

### MiR-210 deficiency affects hepatic lipid metabolism in MASLD mouse model

To elucidate the mechanisms by which miR-210 regulates hepatic lipid accumulation in MASLD, we performed RNA sequencing using liver tissues from WT and miR-210 KO mice fed either CoD or WeD.

We first conducted Gene Set Enrichment Analysis (GSEA) of KEGG, GO and HALLMARK pathways using all detected genes, comparing livers from KO-WeD and WT-WeD mice. The analysis showed significant activation of lipid metabolism-related pathways in KO-WeD mice, including biosynthesis of unsaturated FA, FA metabolism, FA elongation, adipogenesis, and PPARγ signaling pathways (Fig. 4A).

**Fig. 4.**
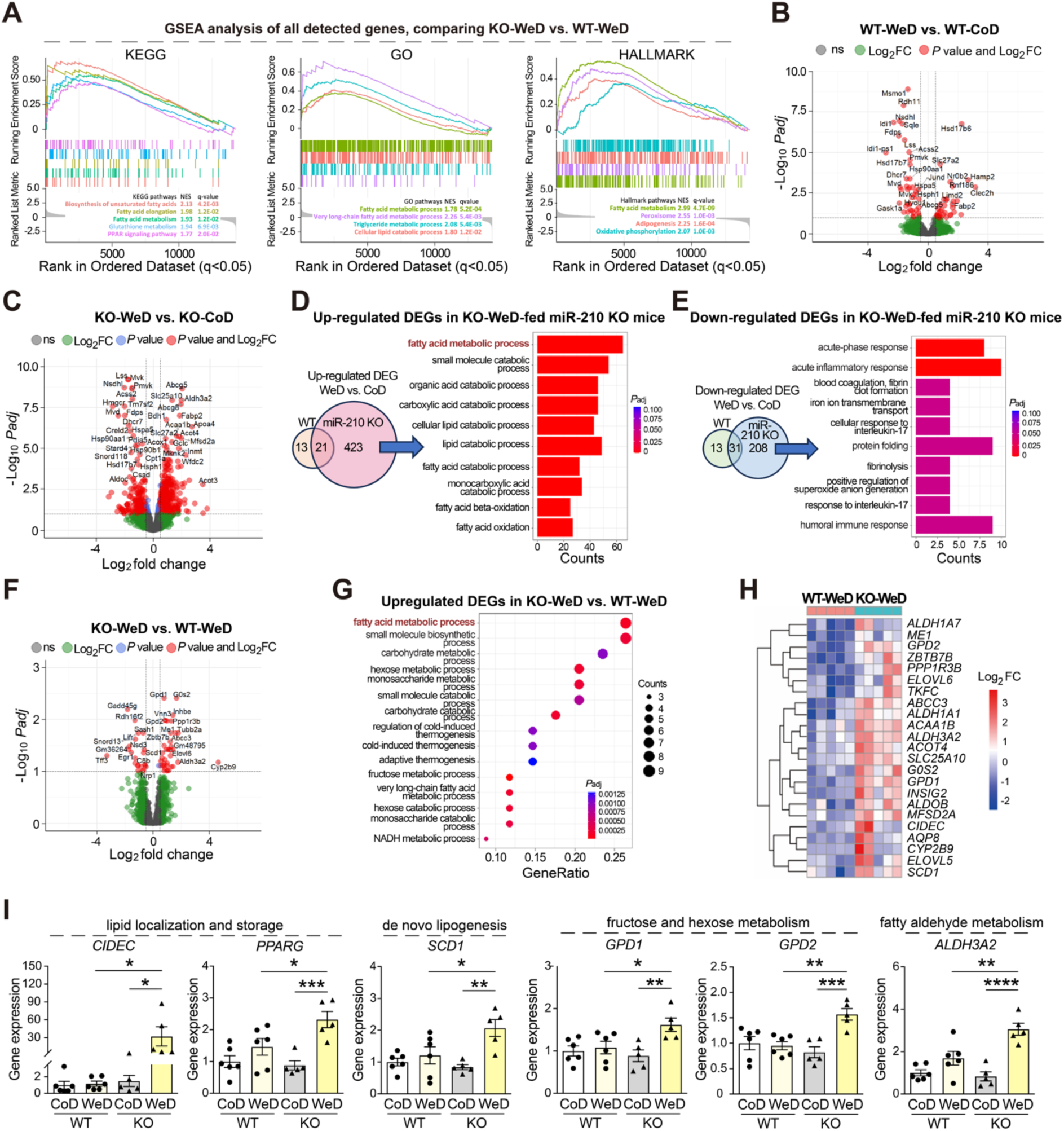
miR-210 deficiency activates hepatic lipid metabolism pathways in MASLD mouse model. Wild-type (WT) and miR-210 knockout (KO) mice were fed a Control (CoD) or Western Diet (WeD) for 10 weeks (n=5-6). RNA sequencing was performed on liver tissues. **(A)** Gene Set Enrichment Analysis (GSEA) of KEGG, GO and HALMARK pathways, comparing livers from WeD-fed miR-210 KO mice (KO-WeD) to WeD-fed WT mice (WT-WeD). Top up-regulated pathways with *q* value < 0.05 are shown. **(B-C)** Volcano plots showing differentially expressed genes (DEGs) in response to WeD in WT **(B)** and miR-210 KO **(C)** mice. Venn diagram illustrating unique and shared DEGs induced by WeD in WT and KO mice, with pathway enrichment of 423 uniquely upregulated **(D)** and 208 uniquely downregulated **(E)** genes in KO-WeD livers. **(F)** Volcano plots showing DEGs between KO-WeD and WT-WeD mouse livers, and **(G)** GO pathway enrichment of the upregulated DEGs. **(H)** Heatmap of key DEGs known to regulate lipid metabolism. **(I)** qPCR validation of candidate miR-210 target genes from the lipid metabolism-related DEGs that contain predicted miR-210 binding sites in their 3’ UTRs. Data are presented as mean ± SEM. Statistical significance was determined using Two-way ANOVA with Bonferroni post hoc test. \**p* < 0.05; \*\**p* < 0.01; \*\*\**p* < 0.001; \*\*\*\**p* < 0.0001. DEGs (red) in volcano plots (B, C and F): | log_2_Fold Change (FC) | > 0.5 and -log_10_ adjusted *p*-value (*P*adj) > 1.0.

We then analyzed differentially expressed genes (DEGs) induced by WeD in both WT and miR-210 KO mice (Fig. 4B - 4C). A greater number of DEGs were identified in miR-210 KO mice than in WT mice following WeD feeding, consistent with the inhibitory role of miR-210 on gene expression. Venn diagrams comparing DEGs from WT-WeD vs WT-CoD with those from KO-WeD vs KO-CoD showed that miR-210 deficiency uniquely upregulated 423 DEGs under WeD conditions. These genes were predominantly enriched in lipid metabolism-related pathways (Fig. 4D). Additionally, 208 genes were uniquely downregulated in KO-WeD livers, with enrichment in pathways such as acute inflammatory response and protein folding (Fig. 4E). Further comparison between KO-WeD and WT-WeD groups revealed that the upregulated DEGs in KO-WeD livers were significantly enriched in FA and carbohydrate metabolism pathways (Fig. 4F-4G).

These findings from both GSEA and DEG pathway analysis suggest that miR-210 deficiency affected lipid metabolism, which is consistent with the phenotype of increased hepatic lipid accumulation in KO-WeD mice.

We then focused on lipid metabolism-related DEGs between KO-WeD and WT-WeD mice, and the fold changes in their gene counts are shown in Fig. 4H. To identify potential direct mRNA target of miR-210, we screened these DEGs for predicted miR-210 binding sites in their 3’ untranslated region (3’ UTR) using RNA22 v2 and miRWalk. Six candidate genes were identified and confirmed to be upregulated in KO-WeD livers by qPCR (Fig. 4I). These included *CIDEC* and *PPARG* involved in lipid localization and storage, and *SCD1* involved in de novo lipogenesis. The upregulation of these genes may directly enhance the hepatic lipid accumulation in KO-WeD mice. Other genes included *GPD1* and *GPD2* involved in fructose and hexose metabolism and *ALDH3A2* involved in fatty aldehyde metabolism. Among these genes, the increase of *CIDEC* gene was remarkably higher (about 20-fold) in miR-210 KO mice on WeD (Fig. 4I).

Overall, these findings indicate that miR-210 deficiency leads to de-repression of lipid metabolism genes, promoting enhanced hepatic lipid accumulation in response to WeD.

### miR-210 inhibits CIDEC expression and lipid accumulation in hypoxic hepatocytes

To further investigate how miR-210 modulates hepatic lipid accumulation, we exposed HepG2 cells to FAs in normoxia and hypoxia. In agreement with the human hepatic spheroids data, hypoxia increased TG levels in HepG2 cells (Fig. S4A), along with expression of *CD36, CIDEC, PPARG* and *SCD1* genes that were involved in FA transport, localization and storage (Fig. S4B). However, the expression of *GPD1, GPD2* and *ALDH3A2* genes remained unchanged (Fig. S4C).

Transfection of hypoxic HepG2 cells with miR-210 mimic significantly reduced intracellular TG levels (Fig. 5A-5B). Notably, miR-210 mimic selectively suppressed *CIDEC* gene expression, without affecting the expression of *PPARG* and *SCD1* genes (Fig. 5C), or *CD36*, *GPD1, GPD2, ALDH3A2,* and *ELOVL6* genes (Fig. S5). The inhibition of CIDEC by miR-210 mimic was further confirmed at the protein level by Western blotting analysis (Fig. 5D).

**Fig. 5.**
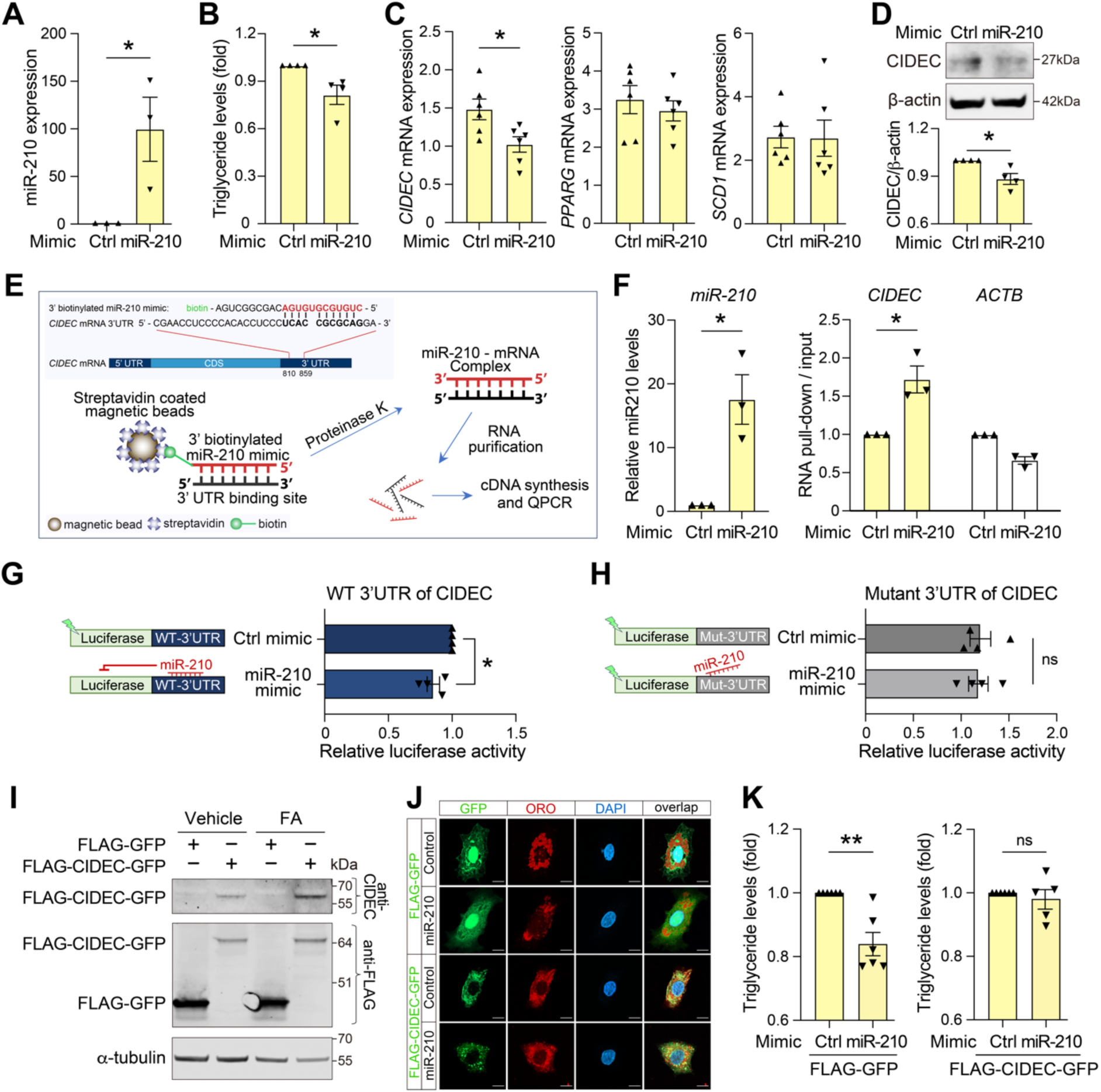
miR-210 inhibits CIDEC expression and lipid accumulation in hypoxic hepatocytes (A-D) HepG2 cells were transfected with 1nM Control mimic (Ctrl) or miR-210 mimic, then treated with fatty acids and cultured under hypoxic conditions. miR-210 expression (**A**), intracellular triglyceride levels (**B**), and gene expression (**C**) were analyzed. (**D**) CIDEC protein expression was assessed by Western blotting. Quantification is shown in the histogram. **(E-F)** RNA pull-down assays. **(E)** Schematic illustration of the RNA pull-down experiment. Sequences of biotin-conjugated miR-210 mimic and the *CIDEC* mRNA 3’UTR region containing miR-210 binding site are shown. HepG2 cells were transfected with biotin-conjugated miR-210 mimic or control mimic, treated with fatty acids under hypoxic conditions. Streptavidin-coated magnetic beads captured biotin-labeled miR-210 and associated mRNAs via its binding sites in 3’ UTR. **(F)** qPCR analysis of miR-210, *CIDEC* and *ACTB* (negative control) mRNAs in the pull-down fraction. **(G–H)** Dual-luciferase reporter assay in HepG2 cells co-transfected with WT **(G)** or mutant (Mut, **H**) *CIDEC* 3′UTR luciferase reporter and control or miR-210 mimic, followed by exposure to fatty acids and hypoxia. **(I)** Western blot validation of FLAG-CIDEC-GFP and FLAG-GFP expression using anti-CIDEC and anti-FLAG antibodies. α-tubulin was used as loading control. **(J)** Representative confocal images demonstrating FLAG-GFP or FLAG-CIDEC-GFP expression (green), lipid droplets stained with Oil Red O (red), and nuclei stained with DAPI (blue). Scale bar: 10 μm. **(K)** Quantification of intracellular triglyceride levels under the indicated conditions. Data are presented as mean ± SEM; n=3-6. Statistical significance was determined by unpaired Student’s t-test test or Mann-Whitney U test. \**p* < 0.05; ** *p* < 0.01. ns: no significant difference.

Taken together, these results suggest that miR-210 regulates lipid accumulation in hypoxic hepatocytes, at least in part, through the downregulation of *CIDEC* gene expression.

### CIDEC identified as a direct target of miR-210

To determine whether CIDEC is a direct target of miR-210, we performed RNA pull-down assays (Fig. 5E). HepG2 cells were transfected with biotin-conjugated miR-210 mimic or control mimic, followed by exposure to FA in hypoxia. Cell lysates were incubated with streptavidin-coated magnetic beads to capture miR-210 and its associated mRNAs via biotin-streptavidin interaction. The pull-down fractions were analyzed by qPCR (Fig. 5F), which revealed significant enrichment of the *CIDEC* mRNA 3’UTR in the miR-210 pull-down, whereas *ACTB* mRNA (negative control) showed no enrichment, indicating direct interaction between miR-210 and the 3’ UTR of *CIDEC*.

To further validate this direct interaction, we performed a dual-luciferase reporter assay using constructs containing either the WT human *CIDEC* 3′UTR or a mutant 3′UTR in which the predicted miR-210 binding site was deleted (Fig. 5G-5H). Co-transfection of HepG2 cells with the WT reporter and miR-210 mimic resulted in a significant reduction in luciferase activity (Fig. 5G). In contrast, the inhibitory effect of miR-210 was abolished when the mutant reporter was used (Fig. 5H), demonstrating that the predicted miR-210 binding site is essential for the interaction and the regulation of *CIDEC* gene by miR-210. Together, the RNA pull-down and luciferase reporter assays confirm that *CIDEC* is a direct downstream target of miR-210 through specific binding to its 3′UTR.

### CIDEC mediates the inhibitory effects of miR-210 on lipid accumulation in hypoxic hepatocytes

To evaluate whether CIDEC mediates the effect of miR-210 on hepatic lipid accumulation, we investigated whether CIDEC overexpression could counteract the lipid-lowering effect of miR-210. Expression of FLAG-CIDEC-GFP and FLAG-GFP control constructs was confirmed by immunoblotting using anti-CIDEC and anti-FLAG antibodies (Fig. 5I). These constructs were co-transfected into HepG2 cells along with miR-210 mimic or control mimic. After treatment with FA and exposure to hypoxia, cells were stained by Oil Red O and analyzed by confocal microscopy (Fig. 5J). Intracellular TG levels were also quantified (Fig. 5K). FLAG-GFP showed diffuse signal all over the cells, whereas FLAG-CIDEC-GFP was predominantly localized around lipid droplets in the cytoplasmic compartment (Fig. 5J). Oil Red O staining and intracellular TG quantification revealed that miR-210 mimic significantly reduced lipid accumulation in cells expressing FLAG-GFP, but this effect was abolished in cells overexpressing FLAG-CIDEC-GFP (Fig. 5J and 5K).

Taken together, these findings establish *CIDEC* as a direct target of miR-210 and highlight its pivotal role in mediating the inhibitory effects of miR-210 on lipid accumulation in hypoxic hepatocytes.

### miR-210 reconstitution inhibits hepatic CIDEC expression and lipid accumulation in miR-210 KO mice fed with Western Diet

Consistent with elevated *CIDEC* mRNA levels in WeD-fed miR-210 KO mice (Fig. 4I), we also detected increased CIDEC protein expression in the livers of these mice (Fig. 6A). To further validate the role of miR-210 in regulating hepatic CIDEC expression and lipid accumulation, miR-210 KO mice fed a WeD were administrated either miR-210 mimic or control mimic via tail vein injection for five days (Fig. 6B). miR-210 mimic specifically suppressed CIDEC mRNA and protein levels in the livers (Fig. 6C-6D), without affecting other candidate genes (Fig. S6A). Moreover, reconstitution of miR-210 led to a significant reduction in hepatic lipid accumulation in WeD-fed miR-210 KO mice, as evidenced by decreased Oil Red O staining (Fig. 6E) and lower intracellular TG levels (Fig. 6F). Interestingly, lipid droplets in the livers of miR-210 mimic-treated mice appeared smaller (Fig. 6E), which is in agreement with previous findings when CIDEC expression is inhibited ^[44]^.

**Fig. 6.**
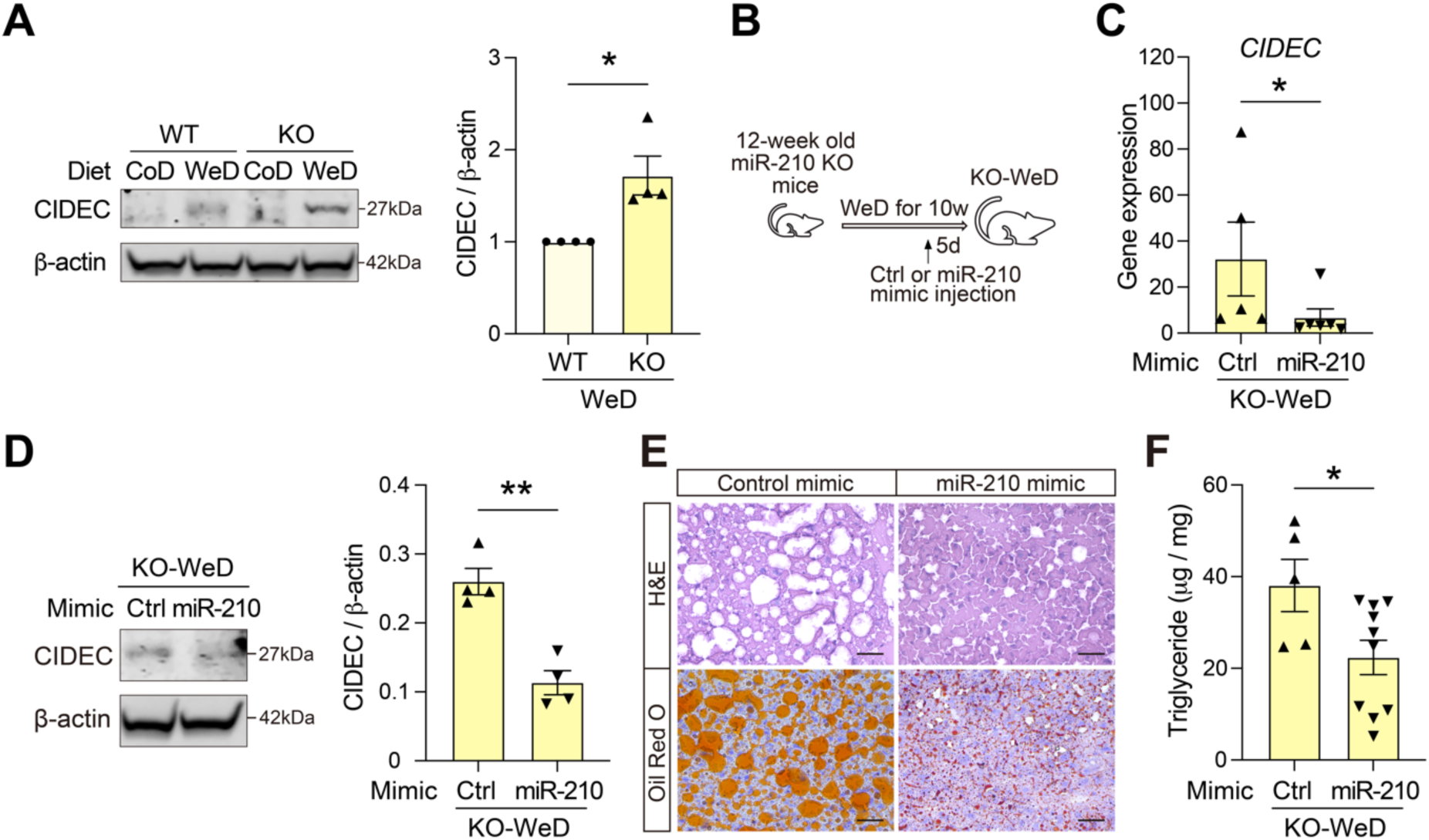
miR-210 reconstitution inhibits hepatic CIDEC expression and lipid accumulation in miR-210 KO mice fed a Western diet. **(A)** Western blot analysis of CIDEC protein expression in liver tissues of wild-type (WT) and miR-210 knockout (KO) mice fed a Control diet (CoD) or Western diet (WeD). ý-actin was used as a loading control. Quantification is shown in histogram. **(B)** Schematic overview of miR-210 mimic or control mimic (Ctrl) administration via tail vein injection in WeD-fed miR-210 KO mice. **(C)** qPCR analysis of hepatic *CIDEC* mRNA levels in WeD-fed miR-210 KO mice (KO-WeD) 5 days after administration with miR-210 mimic or control mimic. **(D)** Western blot analysis of hepatic CIDEC protein levels following mimic treatment. Quantification is shown in histogram. **(E)** Representative images of Hematoxylin & eosin (H&E) and Oil Red O staining of liver sections showing lipid accumulation. Scale bar: 25 μm. **(F)** Quantification of hepatic triglyceride (TG) levels in the indicated groups. Data are presented as mean ± SEM; *n* = 4-10 per group. Statistical analysis was performed using unpaired Student’s t-test or Mann-Whitney U test. \**p* < 0.05; \*\**p* < 0.01.

These findings demonstrate that reconstitution of miR-210 reverses the hepatic phenotype of miR-210 KO mice and confirm that miR-210 suppresses hepatic lipid accumulation, at least in part, by downregulation of CIDEC expression.

### miR-210 suppresses CIDEC and alleviates hepatic steatosis in in vitro and in vivo models of MASLD

To test the therapeutic potential of miR-210 mimic, we employed both human liver spheroids and db/db mouse as models of MASLD. Transfection with miR-210 mimic in FA-treated spheroids specifically reduced *CIDEC* expression (Fig. 7A) and attenuated lipid accumulation (Fig. 7B), without affecting the expression of the other candidate genes (Fig. S6B). *In vivo* treatment (tail vein) with miR-210 mimic alleviated hepatic steatosis in db/db mice (Fig. 7C), shown by reduced Oil Red O staining and smaller lipid droplets in liver sections (Fig. 7D) along with a significant reduction in hepatic TG content (Fig. 7E). Collectively, these results demonstrate that miR-210 suppresses *CIDEC* expression and ameliorates hepatic steatosis, supporting its potential as a therapeutic target for MASLD.

**Fig. 7.**
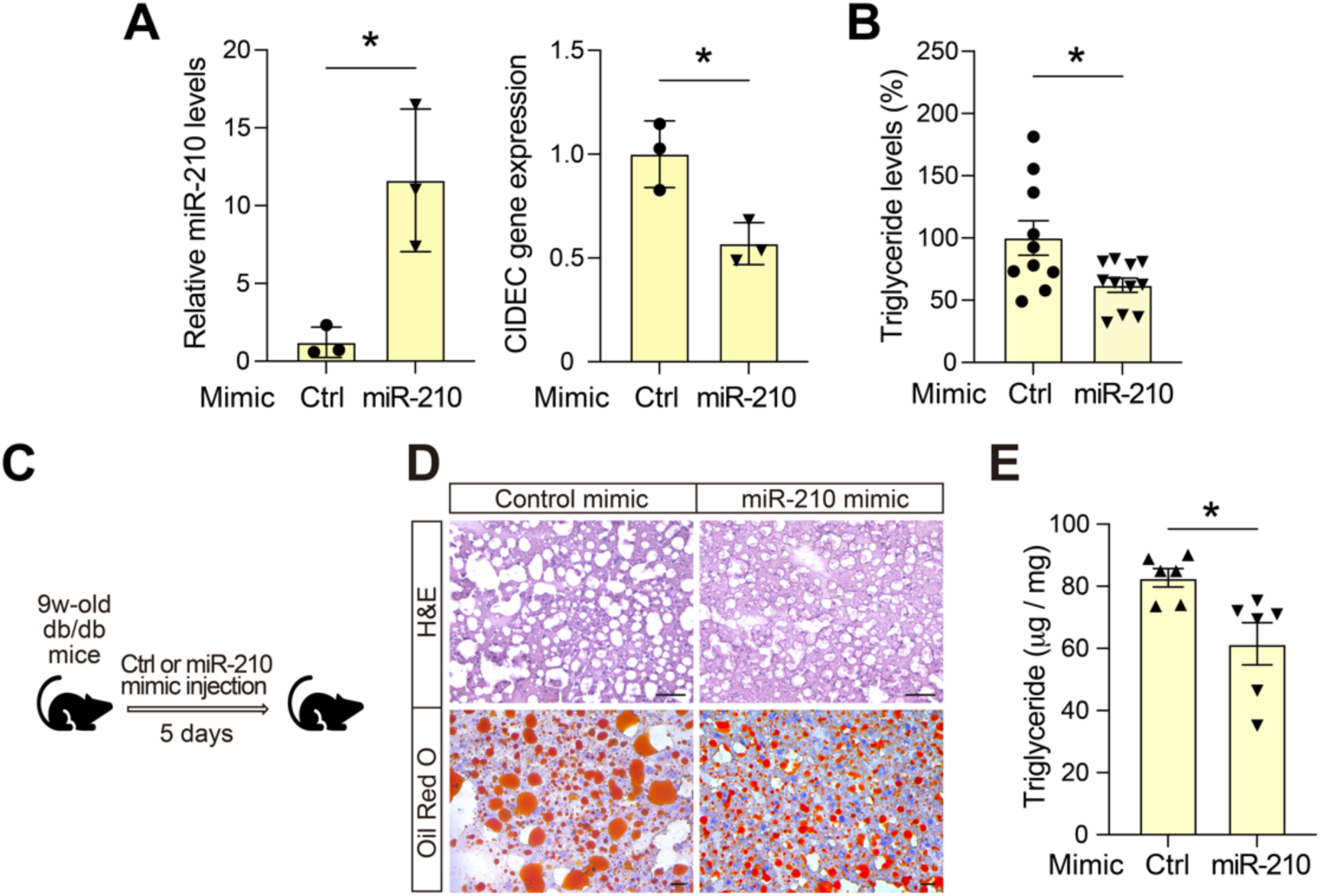
miR-210 inhibits CIDEC expression and alleviates hepatic steatosis in human hepatic spheroids and db/db mice. (A-B) Human hepatic spheroids were transfected with miR-210 mimic or control mimic (Ctrl) and exposed to FA. miR-210 and *CIDEC* mRNA levels were measured by qPCR (**A**, n=3), and lipid accumulation was assessed by AdipoRed staining (**B**, n=10-11). **(C)** Schematic illustration of experiments where db/db mice were administered miR-210 mimic or control mimic via tail vein injection. **(D)** Representative images of Hematoxylin & eosin (H&E) and Oil Red O - stained liver sections from db/db mice treated with control mimic or miR-210 mimic. Scale bar: 25 μm. **(E)** Hepatic triglyceride levels in the indicated groups (n=6). Data are presented as mean ± SEM. Statistical analysis was performed using unpaired Student’s *t*-test. \**p* < 0.05.

## Discussion

This study identifies a previously unrecognized miR-210-CIDEC axis that links FA-induced metabolic stress to hepatic lipid accumulation in MASLD. We demonstrate that while free FAs induce hepatocellular hypoxia, they suppress the canonical hypoxia-responsive induction of miR-210, resulting in de-repression of *CIDEC* gene and exacerbated hepatic triglyceride accumulation. The translational relevance is underscored by reduced serum miR-210 levels in MASLD patients, which correlate negatively with serum TG. Restoring miR-210 through mimic therapy suppresses hepatic CIDEC expression and alleviates steatosis in preclinical models of MASLD, underscoring its promise as a therapeutic strategy.

Hepatic hypoxia is increasingly recognized as a key contributor for MASLD pathogenesis ^[11–15]^. However, the role of HIFs, the principal mediators of cellular adaptation to low oxygen, remains incompletely understood and, in some aspects, controversial ^[11, 17]^. This ambiguity underscores the complexity of hypoxia signaling in MASLD and highlights the need for more specific therapeutic targets. In this study, we identify insufficient miR-210 induction as a critical driver of MASLD and demonstrate that therapeutic administration of miR-210 is sufficient to alleviate hepatic steatosis.

Despite the known hypoxia in MASLD liver, our results demonstrate lower serum miR-210 levels in two independent cohorts of MASLD patients. To our knowledge, it is the first observation regarding serum levels of miR-210 in subjects with MASLD and it is in opposition with subjects with hepatitis where the levels positively correlate with the disease severity ^[45, 46]^. It is important to note that subjects in both MASLD cohorts did not exhibit elevated markers of hepatocellular injury and inflammation, which are likely the source of the high plasma miR-210 levels observed in individuals with hepatitis ^[47]^.

The reverse relationship between TG and miR-210 suggests a repressive effects of FA on miR-210 expression that was confirmed *in vitro*. While FA induced hypoxia in human hepatic spheroids in agreement with previous reports from isolated hepatocytes ^[13],^ this was not followed by an appropriate increase in miR-210 expression. Higher concentration of FA even inhibits miR-210 expression under hypoxic conditions (Fig. 2). Similar repressive effect of FA on hypoxia signaling was reported in the heart ^[48]^. The effect of FA seems to be FA - specific since omega-3 FA have a stimulatory effect on miR-210 levels in cardiomyocytes exposed to hypoxia ^[49]^.

Hypoxia increased intracellular TG levels in both human hepatic spheroids and HepG2 cells, prompting us to further investigate the impact of dysregulated miR-210 expression on lipid accumulation in hepatocytes. Indeed, ablation of miR-210 in mice fed a WeD increased hepatic lipid accumulation, that was reversed by miR-210 reconstitution. Moreover, miR-210 mimic reduced TG levels in hypoxic HepG2 cells, human hepatic spheroids, and livers of db/db mice. Collectively, these findings indicate that impaired miR-210 induction under hypoxia contributes to hepatic lipid accumulation, whereas miR-210 mimic therapy can mitigate it. This is in accordance with the lipid accumulation observed in miR-210-deficient Drosophila retina[22]_._

Transcriptomic analysis revealed a profound modulation of lipid metabolism genes, consistent with the hepatosteatosis phenotype in miR-210 KO mice fed a WeD. We identified CIDEC (also known as FSP27), a lipid droplet-associated protein essential for lipid droplet formation and hepatic triglyceride storage ^[44],^ as a novel miR-210 target mediating these effects. A conserved miR-210 binding site within the 3’UTR of *CIDEC* mRNA was validated by RNA pull-down and reporter assays. miR-210 suppressed the *CIDEC* 3’UTR activity but not that of a mutant lacking the miR-210 binding site. Furthermore, overexpression of CIDEC reversed the inhibitory effects of miR-210 on lipid accumulation in hypoxic hepatocytes.

In WeD-fed miR-210 KO mice, *CIDEC* expression was elevated more than other differentially expressed lipid metabolism genes. miR-210 mimic significantly reduced CIDEC expression in these mice and in human hepatic spheroids, thereby mitigating hepatosteatosis. These findings are consistent with the previous studies demonstrating the critical role of CIDEC in hepatosteatosis in both rodent models and human cohorts ^[50, 51]^. While CIDEC expression is minimal in normal liver, it is markedly upregulated in multiple MASLD models ^[52]^. Both loss-and gain- of function studies support a pathological role of CIDEC in hepatosteatosis ^[50, 51],^ and clinical data further demonstrate CIDEC upregulation in liver from MASLD patients and correlates with steatosis severity ^[50, 53]^.

In summary, our integrated clinical and experimental findings uncover a previously unrecognized mechanism of hepatic steatosis driven by FA-induced dysregulation of miR-210. As illustrated in Figure 8, under physiological conditions, hypoxia induces miR-210, which suppresses CIDEC to limit hepatic lipid accumulation. In contrast, FA overload induces hypoxia but fails to upregulate miR-210, leading to increased CIDEC expression and exacerbated hepatosteatosis. In this context, administration of miR-210 mimic suppresses CIDEC overexpression and attenuates hepatosteatosis. Collectively, these findings identify miR-210-CIDEC axis as a promising therapeutic target for reducing hepatic lipid accumulation and preventing MASLD progression.

**Fig. 8.**
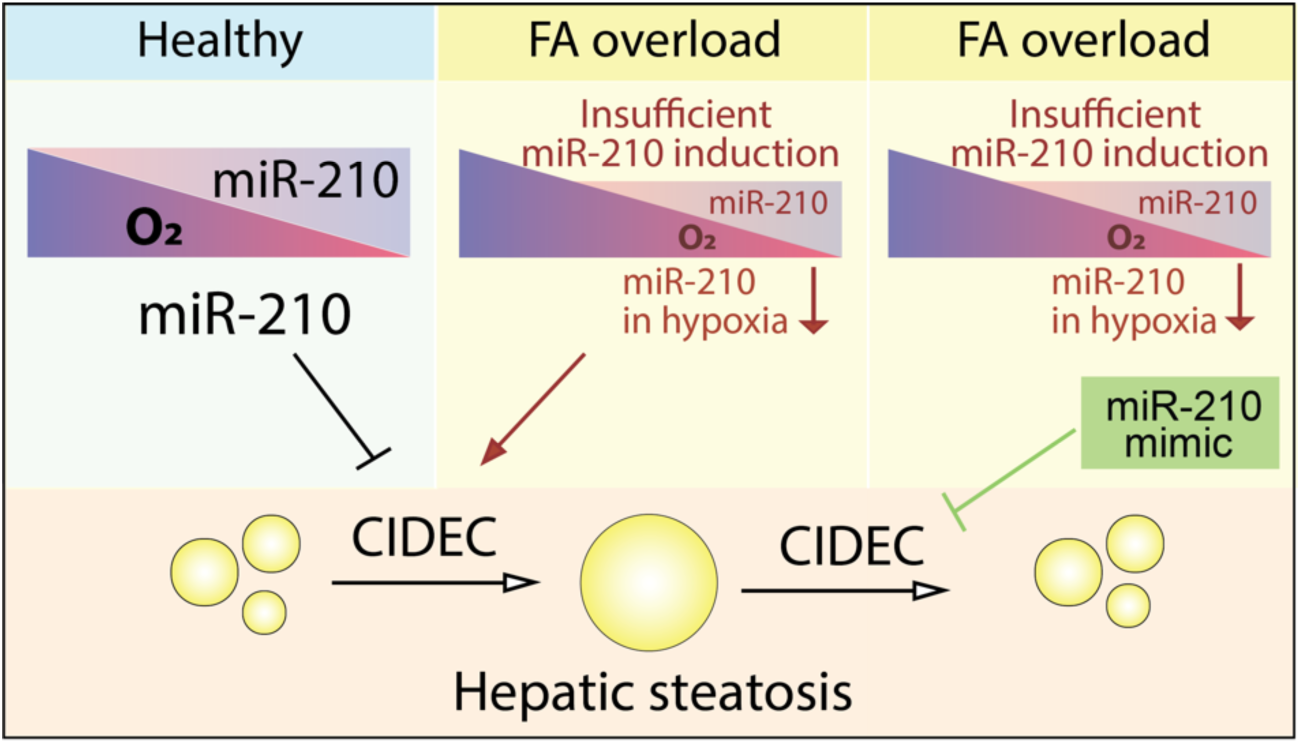
Fatty acid – induced suppression of miR-210 drives hepatic lipid accumulation via CIDEC in metabolic dysfunction-associated steatotic liver disease. Under healthy conditions, hypoxia induces miR-210, which suppresses its target gene CIDEC and thereby limits hepatic steatosis. However, under conditions of FA overload, FA induces hypoxia but inhibits the induction of miR-210, leading to increased CIDEC expression and exacerbated hepatic steatosis. Administration of miR-210 mimic in this setting reduces CIDEC overexpression and attenuates hepatosteatosis. Collectively, these findings identify miR-210 as a promising therapeutic target for reducing hepatic lipid accumulation and preventing MASLD progression.

## Author Contributions

Conceptualization: X.Z. and S-B.C.; Methodology and investigation: B.Y., S.Y., P.C., X.J., Y.I., A.K., J.G., S.N., A.Z. and X.Z. Validation: B.Y., S.Y., P.C., X.J., Y.I., A.K., J.G., Y.L., V.M.L., X.Z. and S-B.C.; Resources: Q.D., N.S., Y.L. V.M.L., X.Z. and S-B.C.; Writing - original draft: B.Y., X.Z. and S-B.C.; Writing review & editing: B.Y., S.Y., P.C., X.J., Y.I., A.K., J.G., S.N., A.Z. Q.D., N.S., Y.L., V.M.L., X.Z. and S-B.C.; Visualization: B.Y. and X.Z.; Supervision: Y.L., V.M.L., X.Z., and S-B.C.; Project administration: X.Z. and S-B.C.; Funding acquisition: Y.L., V.M.L., X.Z., and S-B.C. All authors intellectually commented on and edited the manuscript and approved the final version.

## Declaration of generative AI and AI-assisted technologies in the writing process

During the preparation of this work, the authors used Microsoft Copilot and ChatGPT to assist with grammar checking. After using these tools, the authors reviewed and edited the content as needed and take full responsibility for the content of the publication.

## Data availability statement

The data that support the findings of this study are available from the corresponding author upon reasonable request. RNA sequencing data have been deposited in Gene Expression Omnibus and are accessible through GEO Series accession number GSE313789 (https://www.ncbi.nlm.nih.gov/geo/query/acc.cgi?&acc=GSE313789).

### Abbreviations

CIDE: Cell death–inducing DNA fragmentation factor-α–like effector;
FSP27: fat-specific protein 27;
CoD: Control diet;
FA: Fatty acid;
HIF: Hypoxia-inducible factor;
KO: Knock out;
MASLD: Metabolic dysfunction-associated steatotic liver disease;
NAS: NAFLD Activity Score;
TG: Triglycerides;
WeD: Western Diet;
WT: Wild type;
3’UTR: 3’ untranslated region;
CPM: Counts per million;
DEG: Differentially expressed gene;
GO: Gene ontology;
GSEA: Gene set enrichment analysis.

