## Supplementary material for "HypoxamicroRNA-210 protects against hepatic steatosis by inhibiting CIDEC expression"

**Short title: HypoxamiR-210 prevents hepatic steatosis via CIDEC**

Bo Yan[1,2], Sonia Youhanna[3‡], Peiyin Chen[1‡], Xiuli Jin[1,2], Yuma Iwamura[1,4], Aurino Kemas[3], Jacob Grünler[1], Sampath Narayanan[1], Allan Zhao[3], Qiaolin Deng[3], Norio Suzuki[4], Yiling Li[2†], Volker Martin Lauschke[3,5,6,7], Xiaowei Zheng[1#†], Sergiu-Bogdan Catrina[1,8#†]

[1] Department of Molecular Medicine and Surgery, Karolinska Institutet, Stockholm, Sweden.

[2] Department of Gastroenterology and Hepatology, The First Hospital of China Medical University, Shenyang, Liaoning, China

[3] Department of Physiology and Pharmacology, and Center for Molecular Medicine, Karolinska Institutet and University Hospital, Stockholm, Sweden

[4] Division of Oxygen Biology, Tohoku University Graduate School of Medicine, Sendai, Japan.

[5] Dr Margarete Fischer-Bosch Institute of Clinical Pharmacology, Stuttgart, Germany

[6] University of Tübingen, Tübingen, Germany

[7] Department of Pharmacy, the Second Xiangya Hospital, Central South University, Changsha, China

[8] Center for Diabetes, Academic Specialist Centrum, Stockholm, Sweden

[‡] These authors contributed equally to this work.

[#] Senior authors.

[†] Corresponding authors:

Sergiu-Bogdan Catrina: Department of Molecular Medicine and Surgery, Karolinska Institutet, Stockholm, Sweden. Tel: +46 812367150.

Xiaowei Zheng: Department of Molecular Medicine and Surgery, Karolinska Institutet, Stockholm, Sweden. Tel: +46 735055285..

Yiling Li: Department of Gastroenterology and Hepatology, The First Hospital of China Medical University, Shenyang, Liaoning, China. Tel: +86 13998841476..

### Supplementary methods

#### *Antibodies*

| Name | Citation | Supplier | Cat no. | Clone no. |
|---|---|---|---|---|
| Rabbit anti- pimonidazole antibody | | Hypoxyprobe | PAb2627AP | N/A |
| Rabbit anti- Histone3 antibody | PMID: 37993255 | Abcam | Ab1791 | H3C1 |
| Rabbit anti- Cidec/ FSP27 antibody | PMID: 31097771 | Abcam | ab198204 | N/A |
| Mouse anti-GPD1 antibody | PMID: 33502933 | Santa Crutz | sc376219 | E-7 |
| Rabbit anti- β- actin antibody | PMID: 38114473 | Abcam | ab8227 | N/A |
| Rabbit anti- Flag antibody | PMID: 31511573 | Millpore | F7425 | N/A |
| Mouse anti- α-tubulin antibody | PMID: 35164902 | Abnova | MAB11106 | DMIA |
| IRDye 680RD Goat anti-Rabbit IgG Secondary Antibody | | Li-Cor | 926-68071 | |
| IRDye 800CW Goat anti-Rabbit IgG Secondary Antibody | | Li-Cor | 926-32211 | |
| IRDye 680RD Goat anti- mouse IgG Secondary Antibody | | Li-Cor | 926-68070 | |

#### *Primers*

| Name | Sequence |
|---|---|
| Mouse *ACTB*-F | AAGATCAAGATCATTGCTCCTC |
| Mouse *ACTB*-R | GGACTCATCGTACTCCTG |
| Mouse *CD36*-F | CATTTGCAGGTCTATCTACG |
| Mouse *CD36*-R | CAATGTCTAGCACACCATAAG |
| Mouse *SREBF1*-F | GACATCGAAGACATGCTCCAG |
| Mouse *SREBF1*-R | TCTTTGATCCCAGGCCCAG |
| Mouse F*ASN*-F | GATTCAGGGAGTGGATATTG |
| Mouse *FASN*-R | CATTCAGAATCGTGGCATAG |
| Mouse M*MP9*-F | CTTCCAGTACCAAGACAAAG |
| Mouse *MMP9*-R | ACCTTGTTCACCTCATTTTG |
| Mouse *ACTA2*-F | TGACCCAGATTATGTTTGAGACC |
| Mouse *ACTA2*-R | CAGAGTCCAGCACAATACCAG |
| Mouse *F4/80*-F | CACATCCAGCCAAAGCAG |
| Mouse *F4/80*-R | AACAGCACGACACAGCAG |
| Mouse *TGFB*-F | TGATACGCCTGAGTGGCTGTCT |
| Mouse *TGFB*-R | CACAAGAGCAGTGAGCGCTGAA |
| Mouse *IL1B*-F | GCACGATGCACCTGTACGAT |
| Mouse *IL1B*-R | CACCAAGCTTTTTTGCTGTGAGT |
| Mouse *SCD1*-F | CACCACAAGTTCTCAGAAACAC |
| Mouse *SCD1*-R | GGCTTGTAGTACCTCCTCTG |
| Mouse *ALDH3A2*-F | GTAACAATAAGCTCATCAAACGGG |
| Mouse *ALDH3A2*-R | CTCCAAAGGGCAGAGAATTAACAG |
| Mouse *CIDEC*-F | GAAACTTAAAGACAAGCCCTTCTC |
| Mouse *CIDEC*-R | TCTCTTCTTGCGCTGTTCTG |
| Mouse *PPARG*-F | CCACAAGCATCAAAGTAAGAGAC |
| Mouse *PPARG*-R | TGATCGCACTTTGGTATTCTTGG |
| Mouse *GPD1*-F | AGGAGAAGTTCTGTGAGACGA |
| Mouse *GPD1*-R | GCCACTATATTCTTCAAGGCCC |
| Mouse *GPD2*-F | CCTTCGCCAAGCTCTTCTG |
| Mouse *GPD2*-R | CTCAATGGACTTTCCAGTTCGAG |
| Human *ACTB*-F | AAGATCAAGATCATTGCTCCTC |
| Human *ACTB*-R | ACTCGTCATACTCCTGCT |
| Human *CD36*-F | ATAAGCTCATCAAACGGATG |
| Human *CD36*-R | CCAAATGGGAAAGAGTTGAG |
| Human *ELOVL6*-F | GGATACAGTGTTCATGGTCCT |
| Human *ELOVL6*-R | GAAAGCTTCCTTCATGATGCG |
| Human *ALDH3A2*-F | AAAGAAGCCAACACTAAACC |
| Human *ALDH3A2*-R | TGGTCATTTCGTTAAAGGC |
| Human *CIDEC*-F | CAGAGGAGGTACTACAAACC |
| Human *CIDEC*-R | ATAAGGACGATATVVGAAGAG |
| Human *PPARG*-F | TGGAGCCTTAAAGAATGTAGTG |
| Human *PPARG*-R | TATCATCTCCATGAGTCCCAG |
| Human *SCD1*-F | ATGTTGAAGTGAGAAGAGGG |
| Human *SCD1*-R | CAACATGATTTCGGGAGATAG |
| Human *GPD1*-F | ATGTATGTGGAGAAGGGTTGG |
| Human *GPD1*-R | GACACAGATGTTGCTAGGAG |
| Human *GPD2*-F | CCGACCTGTCTGAAGATACTG |
| Human *GPD2*-R | AGTCTGGATGGGCTAAGCTG |

| | |
|---|---|
| Human *CIDEC* 3'UTR-F | AGTATATTCGGTGCTCTTCG |
| Human *CIDEC* 3'UTR-R | TTAGCACAAATGCATAAGCC |

#### *Other materials*

| | | |
|---|---|---|
| miRNeasy Serum/ Plasma Advanced kit | Qiagen | 217204 |
| miRcute Plus miRNA First-Strand cDNA kit | TianGen Biotech (Beijing) | KR211 |
| miRcute Plus miRNA qPCR kit (SYBR Green) | TianGen Biotech (Beijing) | FP411 |
| Oil red O powder | Sigma | O0625 |
| Glycerol mounting medium | DAKO | C0563 |
| TG quantification colorimetric / fluorometric kit | Sigma | MAK266 |
| Hypoxyprobe-1 Omni kit | Hypoxyprobe Inc | |
| Pierce BCA protein assay kit | Thermofisher | 23225 |
| Halt protease and phosphatase cocktail | Thermofisher | 78443 |
| Quant-iT dsDNA assay kits | Thermofisher | Q33120 |
| 4× Blot LDS loading buffer | Invitrogen | B0007 |
| SurePAGE (Bis-Tris, 10×8, 4-20%) | GenScript | M00656 |
| 20×MOPS running buffer | Invitrogen | NP0001 |
| SeeBlue Plus2 pre-strained protein standard | Invitrogen | LC5925 |
| Nupage antioxidant | Invitrogen | NP0005 |
| Immun-Blot Low Fluorescence PVDF Membrane | Bio-Rad | 1620264 |
| 20× transfer buffer | Invitrogen | NP0006 |
| Intercept blocking buffer (TBS) | Li-Cor | 927-60001 |
| NewBlot PVDF stripping buffer | Li-Cor | 928-40032 |
| Direct-zol RNA mini/micro preps kits | Zymo | R2050/2060 |
| TaqMan Advanced miRNA cDNA synthesis kits | Thermofisher | A28007 |
| Taqman Advanced miRNA Assay | Thermofisher | A25576 |
| Taqman Fast Advanced Master mix | Thermofisher | 4444963 |
| OCT | Tissue-Tek | 4853 |
| Limonene Mounting Medium | Abcam | ab104141 |
| Nucleospin miRNA kits | MACHEREY-NAGEL | 740971 |
| RNAlater solution | Thermofisher | AM7021 |
| High-Capacity cDNA reverse transcription kits | Thermofisher | 4368814 |
| RNase inhibitor | Thermofisher | N8080119 |
| Maxima H minus reverse transcriptase | Thermofisher | EP0752 |
| 5×Maxima RT buffer | Thermofisher | EP0752 |
| 25mM dNTP | NEB | N0447L |
| 100uM TSO | E5V7NEXT | |
| 10uM barcoded oligo dT | E5V7NEXT | |
| Exonuclease | NEB | M0293S |
| 10×ExoI buffer | NEB | |
| 2×KAPA HiFi Ready Mix | Kapa biosystems | KR0370 |
| 10 µM pre-amp primer | SINGV6 | |
| Quanti-iT PicoGreen dsDNA assay kit | Life Technologies | P7589 |
| High Sensitivity DNA Analysis kit | Aligent | 5067-4626 |
| NEBNext MLtra II FS DNA Library Prep kit for Illumina | NEB | E7805S |
| Sera-Mag Speedbead carboxylate-modified [E3] magnetic particles | Cytiva | 65152105050250 |
| Lipofectamine RNAiMAX | Thermofisher | 13778075 |
| Streptavidin coated magnetic beads | Thermofisher | 88816 |
| Superscript IV First Strand cDNA Synthesis kit | Thermofisher | 18091050 |
| Lipofectamine 3000 Transfection Reagent | Thermofisher | L3000015 |
| Isopropanol | Histolab | -/02155 |
| Penicillin (10000 U/ ml) -Streptomycin (10000 µg/ ml) | Gibco | 15140122 |
| Sodium Pyruvate (100mM) | Gibco | 11360070 |
| Phosphate-buffered saline | Gibco | 14190094 |
| Dimethyl sulfoxide, DMSO | Sigma | D8418 |
| Palmitic acid | Sigma | P0500 |
| Oleic acid | Sigma | O1083 |
| Hematoxylin | Histolab | 01820 |
| 4% formaldehyde | Histolab | 02176 |
| IGEPAL CA-630 | Sigma | 56741 |
| DL-Dithiothreitol | Sigma | D0632 |
| 10% formalin | Sigma | TH501320 |
| PBS tablets | Sigma | P4417 |
| Eosin | Histolab | 01650 |
| Absolute ethanol | Histolab | 01399 |
| 95% ethanol | Histolab | 01396 |
| 70% ethanol | Histolab | 01370 |
| Sodium chloride (NaCl) | Sigma | S9888 |
| Potassium chloride (KCl) | Sigma | 60128 |
| Magnesium chloride ($MgCl_2$) | Sigma | M0250 |
| Glycine | Sigma | G7126 |

| | | |
|---|---|---|
| 4% paraformaldehyde | Histolab | 123630 |
| Ethylene Diamine Tetraacetic Acid | Sigma | ED2SC |
| Trizma base | Sigma | 252859 |
| Sodium dodecyl sulfate | Sigma | L3771 |
| Fetal bovine serum | Gibco | A5256801 |
| Dulbecco's Modified Eagle Medium | Gibco | 61965026 |
| 0.05% trypsin | Gibco | 25300054 |
| T75 Flask culture plate | Sarstedt | 9023511 |
| Microtome N35 blade | pfmmedical | 207500006 |
| Cryomold | Tissue-Tek | 62534 |
| Superforest plus slides | Epradia | J1800AMNZ |
| Cover slides | Menzel | 11911998 |
| Stainless steel beads | Qiagen | 69990 |
| Opti-MEM I Reduced Serum Medium | Gibco | 31985062 |
| RNase inhibitor | Thermofisher | N8080119 |
| Yeast tRNA | Sigma | R8759 |
| Proteinase K | Invitrogen | AM2546 |

#### Supplementary tables

***Table S1. Clinical characteristics of MASLD patients and healthy controls***

| | Control | MASLD | *P* value |
|---|---|---|---|
| Age | 61.27 ± 1.60 | 59.24 ± 0.68 | 0.26 |
| Gender | | | |
| Male | 4 | 5 | |
| Female | 7 | 16 | |
| Body weight (kg) | 62.00 (50.00-65.00) | 65.80 (61.60-68.00) | 0.06 |
| *Metabolic parameters* | | | |
| BMI (kg/m$^2$) | 22.16 ± 0.98 | 25.43 ± 0.58 | 0.008 |
| Fasting glucose (mmol/L) | 4.98 (4.70-5.02) | 5.54 (5.17-6.04) | < 0.001 |
| TG (mmol/L) | 0.93 ± 0.12 | 2.08 ± 0.22 | < 0.001 |
| TC (mmol/L) | 5.12 ± 0.40 | 5.17 ± 0.18 | 0.89 |
| HDL (mmol/L) | 1.38 (1.19-1.98) | 1.18 (1.02-1.46) | 0.02 |
| LDL (mmol/L) | 3.07 ± 0.37 | 3.23 ± 0.17 | 0.23 |
| *Clinical parameters* | | | |
| Albumin (g/L) | 44.19 ±0.58 | 44.60 ± 0.51 | 0.63 |
| AST (U/L) | 22.00 (22.00-46.00) | 29.00 (22.00-39.50) | 0.15 |
| ALT (U/L) | 26.91 ± 6.37 | 34.81 ± 4.80 | 0.24 |
| ALP (U/L) | 79.00 (69.00-99.00) | 94.00 (70.00-110.00) | 0.16 |
| GGT (U/L) | 13.00 (10.00-18.00) | 29.00 (18.00-52.50) | 0.005 |
| Platelet ($10^9$ /L) | 221.4 ± 23.58 | 220.8 ± 12.47 | 0.98 |
| *Fibrotouch parameters* | | | |
| Fatty decay (dB/m) | | 277.00 ± 3.97 | |
| Mild (%) | | 23.81 | |
| Middle (%) | | 61.90 | |
| Severe (%) | | 14.29 | |
| Hepatic elasticity (kPa) | | 7.50 (5.70-8.95) | |
| F0-F1 (%) | | 80.95 | |
| F2-F3 (%) | | 14.29 | |
| F4 (%) | | 4.76 | |

Abbreviations: BMI: body mass index; TG: triglyceride; TC: total cholesterol; HDL: high-density lipoprotein; LDL: low-density lipoprotein; AST: aspartate aminotransferase; ALT: alanine aminotransferase; ALP: alkaline phosphatase; GGT: gamma-glutamyl transferase.

*Table S2. Correlation analysis between serum miR-210 levels and clinical parameters*

| | r | *P* value |
|---|---|---|
| Age | -0.074 | 0.69 |
| Body weight | -0.29 | 0.15 |
| BMI ($kg/m^2$) | -0.32 | 0.11 |
| Fasting glucose (mmol/L) | -0.29 | 0.11 |
| TG (mmol/L) | -0.37 | 0.040 |
| TC (mmol/L) | -0.029 | 0.87 |
| HDL (mmol/L) | 0.52 | 0.0022 |
| LDL (mmol/L) | -0.12 | 0.51 |
| AST (U/L) | -0.088 | 0.63 |
| ALT (U/L) | 0.0087 | 0.96 |
| ALP (U/L) | -0.015 | 0.93 |
| GGT (U/L) | -0.068 | 0.71 |
| Noninvasive risk predictive model of steatosis and fibrosis | | |
| HSI | 0.44 | 0.10 |
| TyG | -0.52 | 0.0021 |
| FIB-4 | -0.048 | 0.84 |
| NFS | 0.50 | 0.05 |
| Fibrotouch parameters | | |
| Fatty decay (dB/m) | 0.34 | 0.14 |
| Hepatic elasticity (kPa) | 0.53 | 0.014 |

Abbreviations: BMI: body mass index; TG: triglyceride; TC: total cholesterol; HDL: high-density lipoprotein; LDL: low-density lipoprotein; AST: aspartate aminotransferase; ALT: alanine aminotransferase; ALP: alkaline phosphatase; GGT: gamma-glutamyl transferase; HIS: hepatic steatosis index; TyG: triglyceride glucose index; FIB-4: Fibrosis-4 index; NFS: non-alcoholic fatty liver disease (NAFLD) fibrosis score.

#### Supplementary figures and figure legends

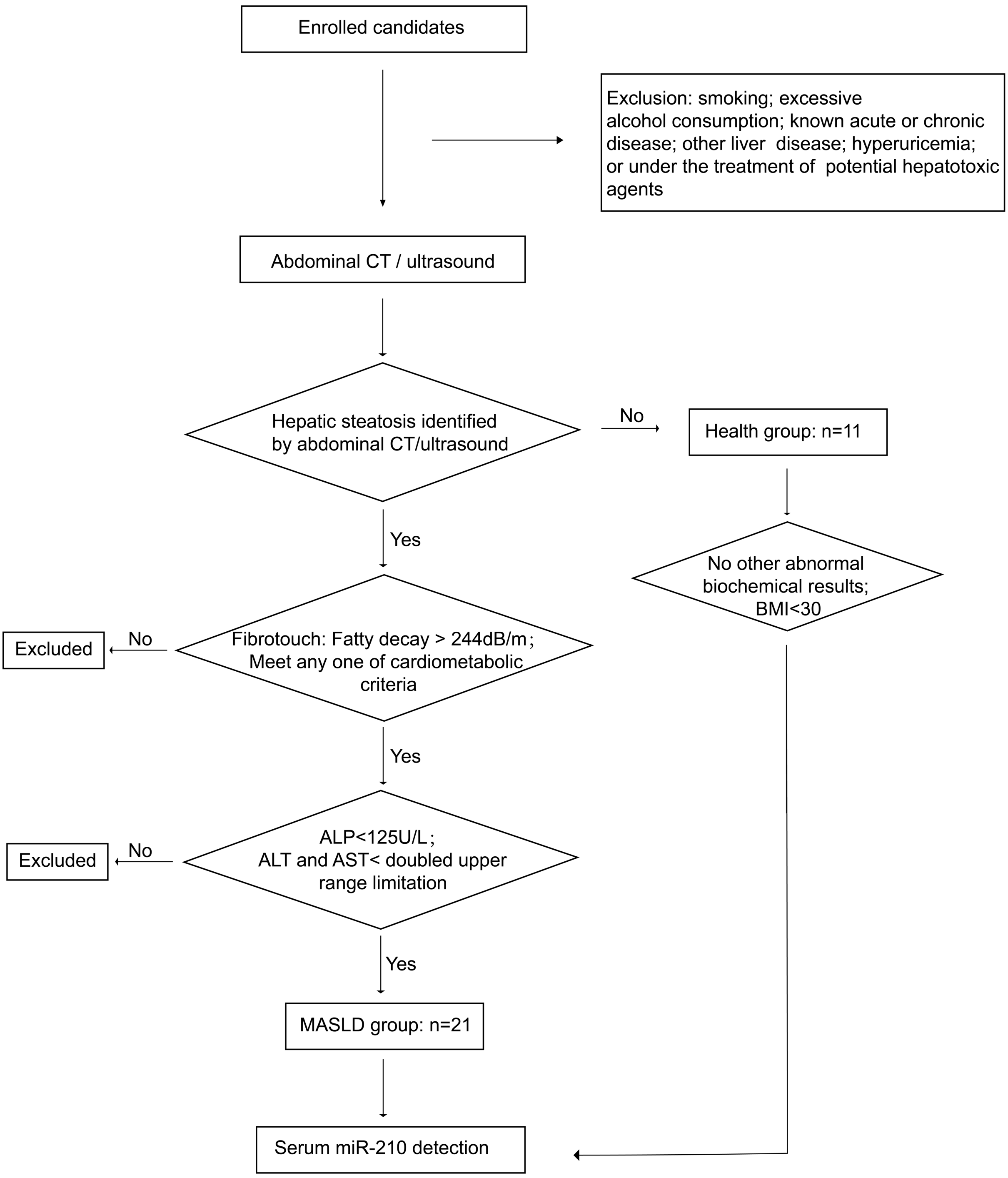

***Fig. S1 (related to Fig. 1)***

Flow chart demonstrating the recruitment of MASLD patients and healthy controls for collection of serum samples.

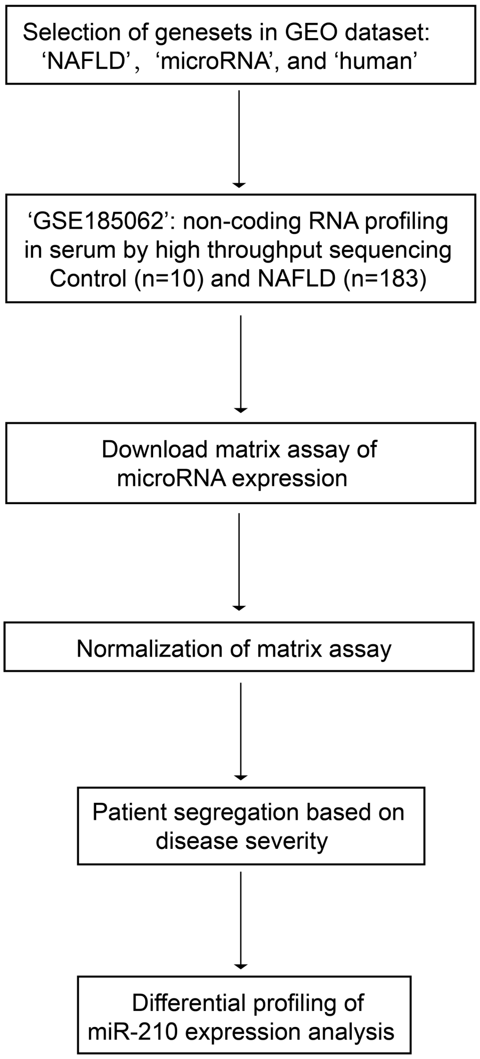

***Fig. S2 (related to Fig. 1)***

Flow chart illustrating the analysis of miR-210 counts in NCBI dataset GSE185062 which includes non-coding RNA sequencing data from the serum of 183 individuals with biopsy-confirmed hepatic steatosis and 10 healthy controls.

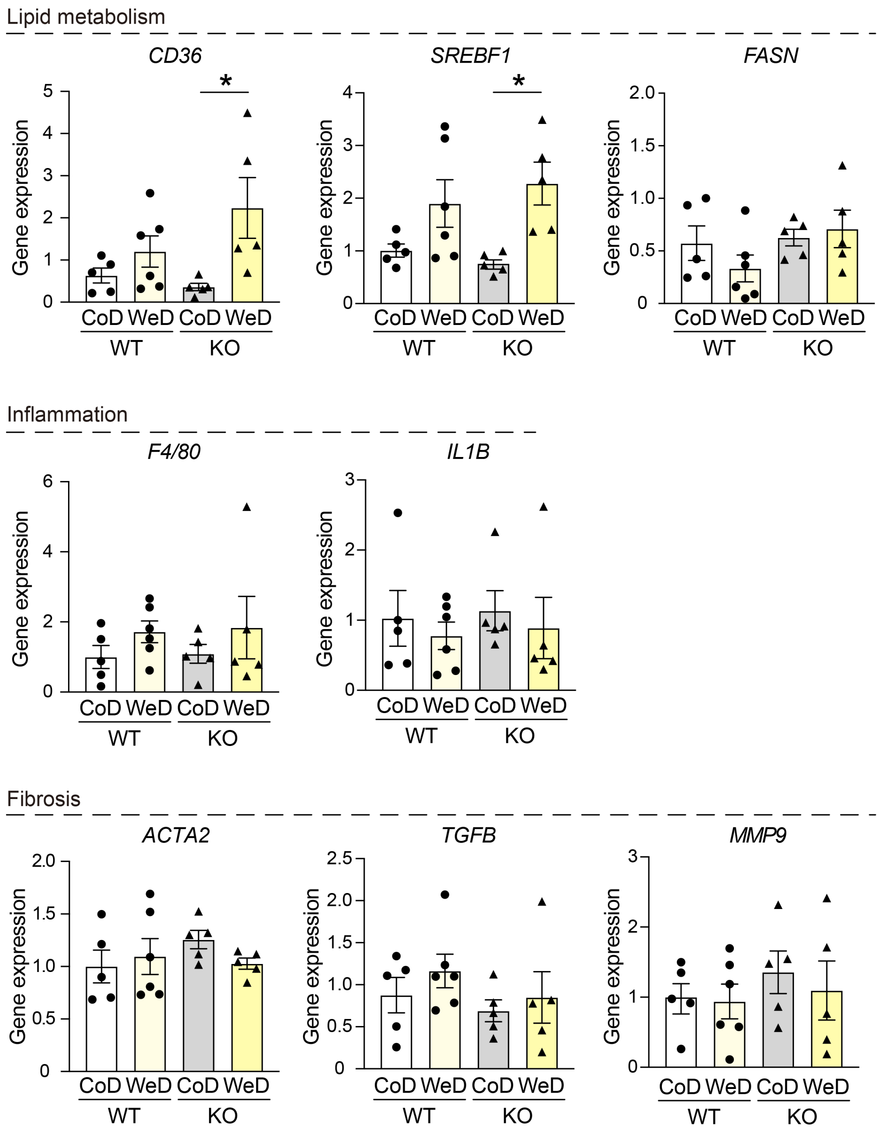

*Fig. S3 (related to Fig. 3)*

**miR-210 deficiency activates lipid metabolism genes, but not inflammation- or fibrosis-related genes in liver from mice on WeD**

Wild-type (WT) and miR-210 knockout (KO) mice were fed a Control (CoD) or Western Diet (WeD) for 10 weeks (n=5-6). qRT-PCR was performed to analyze the relative gene expression of lipid metabolism-, inflammation- and fibrosis-related genes in liver tissue. Data are presented as mean ± SEM. Statistical significance was determined using Two-way ANOVA with Bonferroni post hoc test. *$p < 0.05$.

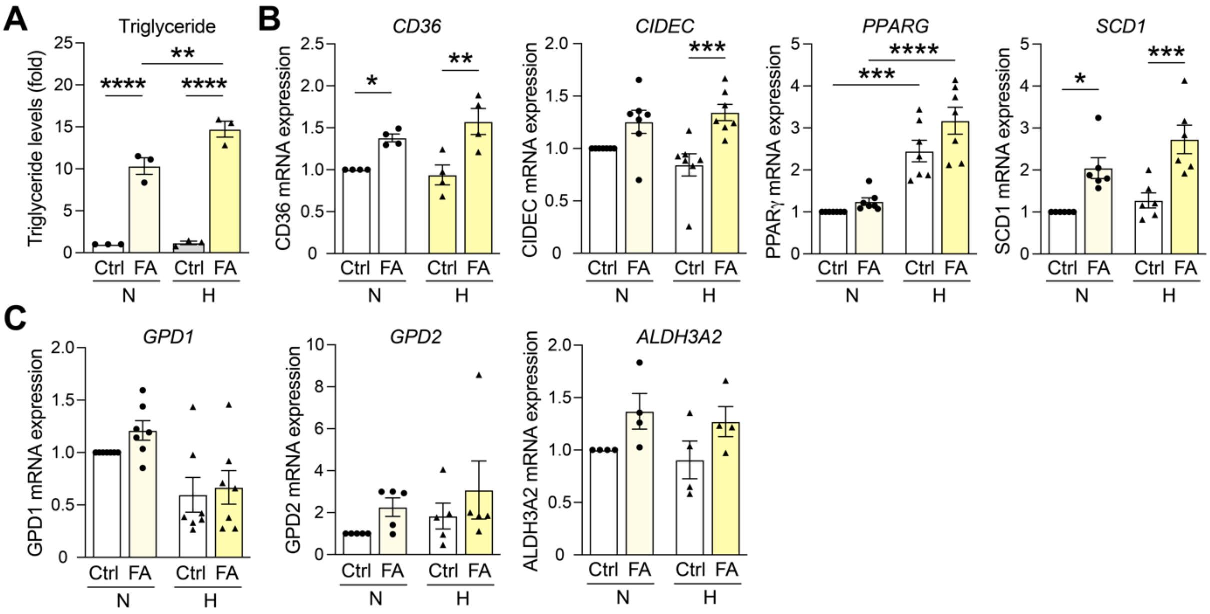

*Fig. S4 (related to Fig. 5)*

**Hypoxia increases triglyceride levels and affects lipid metabolism in HepG2 cells**

Triglyceride levels (**A**) and gene expression (**B-C**) were analyzed in HepG2 hepatocytes treated with fatty acids (FA) and cultured under normoxic (N) or hypoxic (H) conditions. Data are presented as mean ± SEM; n=3-6. Statistical significance was determined using Two-way ANOVA with Bonferroni post hoc test. *$p < 0.05$; ** $p < 0.01$; ***$p < 0.001$; ****$p < 0.0001$.

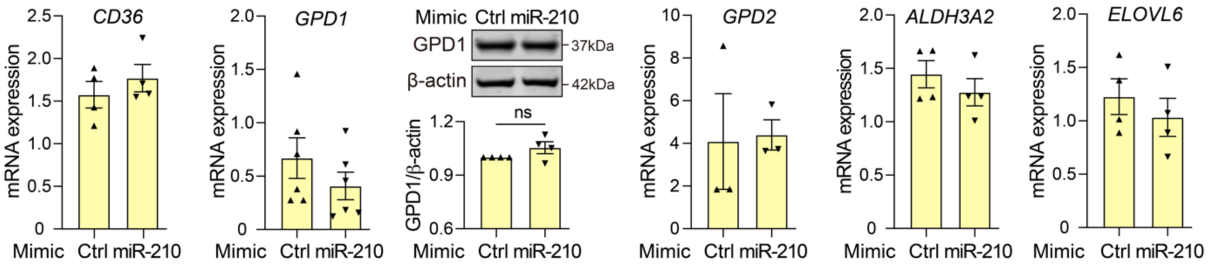

***Fig. S5 (related to Fig. 5)***

**miR-210 did not affect *CD36*, *GPD1*, *GPD2*, *ALDH3A2* and *ELOVL6* gene expression in HepG2 cells**

HepG2 cells were transfected with 1nM Control mimic (Ctrl) or miR-210 mimic, then treated with fatty acids and cultured under hypoxic conditions. Gene expression of *CD36*, *GPD1, GPD2, ALDH3A2* and *ELOVL6* were analyzed by qRT-PCR. GPD1 protein expression was assessed by Western blotting. Quantification is shown in the histogram. Data are presented as mean ± SEM; n=3-6. Statistical significance was determined using unpaired Student's t-test or Mann-Whitney U test.

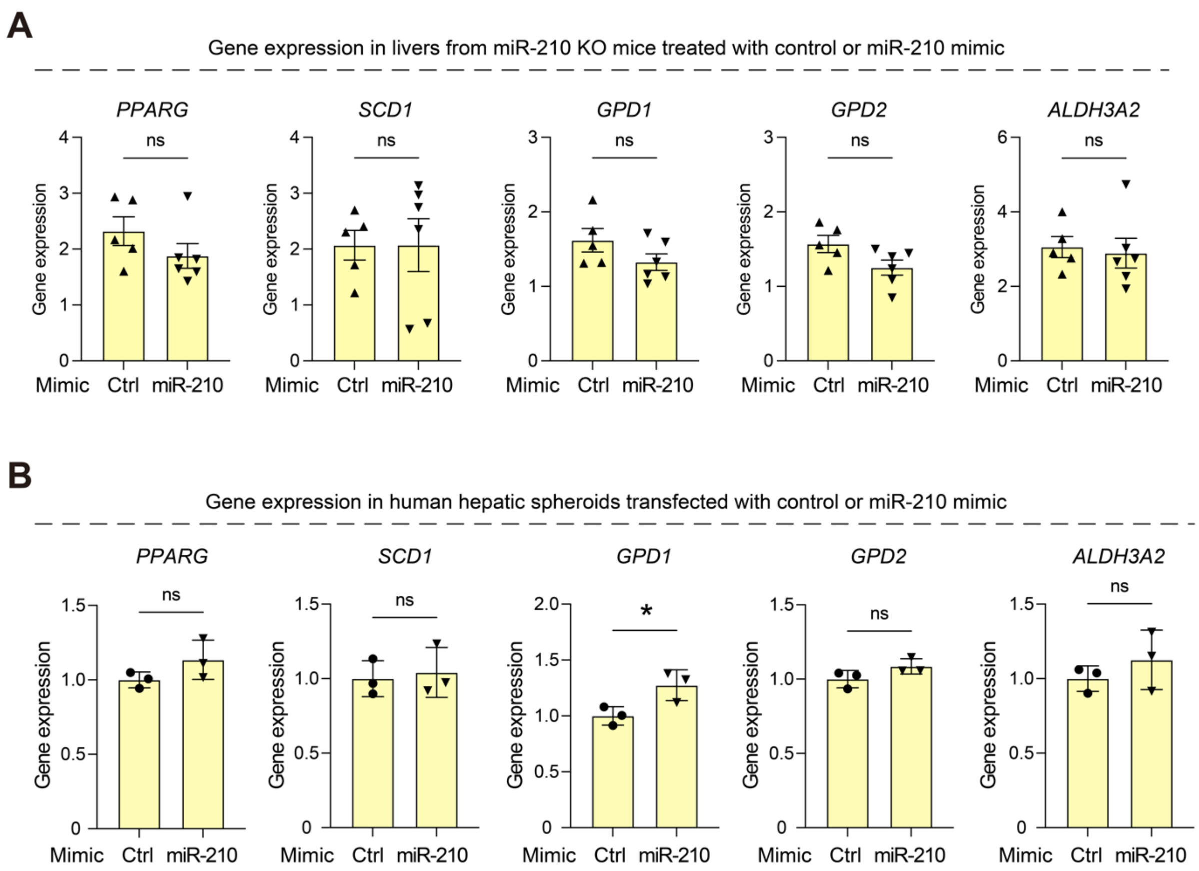

*Fig. S6 (related to Fig. 6 and 7)*

**miR-210 did not repress *PPARG*, *SCD1*, *GPD1, GPD2* or *ALDH3A2* gene expression**

**(A)** qRT-PCR analysis of hepatic gene expression in WeD-fed miR-210 KO mice 5 days after the administration with miR-210 mimic or control mimic (Ctrl) (n=5-6). **(B)** Human hepatic spheroids were transfected with miR-210 mimic or control mimic (Ctrl) and exposed to FA. Expression of indicated genes were measured by qRT-PCR (n=3). Data are presented as mean ± SEM. Statistical significance was determined using unpaired Student's t-test or Mann-Whitney U test. *$p < 0.05$; ns: no significance.
